# L1CAMxCD3 bispecific antibodies exert potent anti-tumor effects in preclinical pancreatic cancer models considering the complex tumor stroma

**DOI:** 10.64898/2026.08.10.743835

**Authors:** Anna Maxi Wandmacher, Annika Brauer, Charlotte Kayser, Caj Stach, Jakob Werner, Silje Beckinger, Tina Daunke, Leon Baumann, Benjamin Heckelmann, Ashinikumar Hidam, Olha Lapshyna, Daniela Wesch, Anne-Sophie Mehdorn, Christoph Röcken, Rüdiger Braun, Flavio Mehli, Anne Schmidt, Gunther Spohn, Susanne Sebens

## Abstract

Pancreatic ductal adenocarcinoma (PDAC) is characterized by an immunosuppressive tumor microenvironment (TME) with pancreatic myofibroblasts (PMF) and macrophages being two prominent cell populations essentially impairing tumor responses to (immuno)therapies. L1 cell adhesion molecule (L1CAM) is upregulated in PDAC cells in primary and metastatic tissues and associated with tumor progression and therapy resistance. Using L1CAM as tumor-associated antigen, two bispecific antibodies (bsAB) targeting L1CAM and CD3 were developed in the IgG-(L)-ScFv format and their anti-tumorigenic activity was investigated in different preclinical PDAC models. In 2D models, both L1-bsAB exerted L1CAM-specific anti-PDAC cell activity when co-cultured with activated CD8+ T cells. Strong anti-PDAC cell effects along with elevated release of T cell effector molecules were also observed upon co-culture with peripheral blood mononuclear cells (PMBC) from healthy donors and PDAC patients. Of note, both L1-bsAB were also effective in 3D PDAC cell spheroids and neither impaired by PMF nor macrophages. Finally, application of L1-bsAB on organotypic tissue slice cultures from PDAC tissues comprising the entire complex TME also induced PDAC cell apoptosis and release of T cell effector molecules. Overall, our results highlight relevant anti-PDAC cell activity of L1-bsAB in immunosuppressive contexts supporting their potential as immunotherapeutic strategy for PDAC.

## Introduction

The incidence of pancreatic ductal adenocarcinoma (PDAC) is rising, whereas only incremental improvements in therapeutic strategies have been achieved lately. The disease is commonly diagnosed at metastasized, non-curative stages due to unspecific symptoms and pronounced therapy resistance resulting in the dismal prognosis and high PDAC-related mortality.^1^ While the advent of RAS inhibitors sparks hope for PDAC patients, several immunotherapeutic strategies have failed to convey significant clinical benefit in PDAC patients.^2–4^ One key factor that can reduce the effectiveness of (immunotherapeutic) strategies is the pronounced tumor stroma of PDAC. Here, stromal cell populations, including pancreatic myofibroblasts (PMF) or macrophages, essentially contribute to an immunosuppressed tumor microenvironment (TME). Macrophages are highly plastic and can thereby exhibit different phenotypes. M1-macrophages are associated with pro-inflammatory activity promoting T cell effector functions, while M2- macrophages are involved in promotion of extracellular matrix (ECM) deposition and secretion of anti-inflammatory cytokines, which can impair the effector phenotype of T cells. With progression of the disease, M2- or alternatively activated macrophages are increasingly enriched in the PDAC TME.^5^ Thus, there is an urgent clinical need for novel therapeutic strategies for PDAC treatment that can overcome these stromal hurdles.

Development of bispecific antibodies, especially T cell engagers (TCE), for the treatment of solid tumors has gained momentum fueled by successful clinical application in hematologic malignancies. TCE mediate tumor-directed killing by direct activation of T cells via simultaneous binding to a surface antigen on target cells and T cells independent of antigen presentation by HLA molecules or the presence of a specific T cell receptor (TCR). Activated T cell populations subsequently kill target cells via release of pore-forming (Perforin) and apoptosis-inducing proteases (Granzymes).^6^ Simultaneously, secreted cytokines amplify the immune response and maintain anti-tumor immune activity. In comparison to adoptive T cell-based strategies such as chimeric antigen receptor (CAR) T-cell therapy, TCE can be applied as off-the-shelf therapeutics with lower manufacturing costs and turn-around times. Additionally, clinical application has a lower risk of high-grade cytokine release syndrome and neurotoxicity.^7^ Development of TCE for treatment of solid tumors has been challenging due to the lack of appropriate tumor specific surface antigens and the presence of an immuno-suppressed TME inhibiting invasion and effector function of T cells. Lately, the approval of Tebentafusp for treatment of uveal melanoma and Tarlatamab for treatment of small cell lung cancer in the U.S. and EU have proven that these obstacles can be overcome successfully.^8^

The neural cell adhesion molecule L1 (L1CAM, CD171) is a glycosylated Type-I transmembrane protein that belongs to the immunoglobulin superfamily. L1CAM was first described as a neuronal cell adhesion molecule involved in neuronal development during embryogenesis. In human adults, it is physiologically expressed at moderate levels in brain, nerve tissue and renal collecting ducts. Of note, strongly elevated protein levels can be detected in up to 80% of primary PDAC specimen as well as in 100% PDAC lymph node and liver metastases.^9^ Preclinical studies have identified multiple tumor-promoting functions of L1CAM, including induction of chemotherapy resistance, tumorigenicity, metastasis formation and immune evasion.^10–15^ Altogether, these findings underscore that L1CAM is an attractive target for an immunotherapeutic strategy of PDAC. Published data on L1CAM-directed antibody-based therapeutics suggest efficacy and good tolerability in animal models of solid cancers.^16–18^ To date, promising preclinical data on targeting L1CAM with monoclonal antibodies or CAR-T cell therapy in solid tumors have not been translated into clinical benefit. Deploying TCE represents a valid therapeutic strategy to improve clinical efficacy of drugs targeting L1CAM circumventing tedious and costly generation of adoptive T cells while maximizing anti-tumor efficacy by local T cell activation.

Thus, this study evaluated the therapeutic potential of two novel L1CAMxCD3 bispecific antibodies (L1-bsAB1 and L1-bsAB2), designed in the IgG-(L)-ScFv format and characterized by high affinity to L1CAM, in different preclinical PDAC model systems accounting for the immunosuppressive TME in PDAC. Our data demonstrate potent preclinical efficacy of L1-bsAB1 and L1-bsAB2 in 2D PDAC models, in 3D spheroids in the absence and presence of different stromal cell populations, i.e. PMF or macrophages, as well as in organotypic slice cultures (OTSC) with a sustained TME obtained from resected PDAC specimens.

## Results

### Characterization of L1-bsAB

L1-bsAB1 and L1bsAB2 were designed in an IgG (L) scFv format, where humanized IgG1 antibodies directed against L1CAM were fused at the C-termini of their light chains to two different scFvs against CD3. In L1-bsAB1, an anti-L1CAM IgG was fused to a scFv derived from the humanized antibody OKT3, whereas in L1-bsAB2 a similar anti-L1CAM IgG was fused to a scFv derived from a humanized version of the SP34 antibody. ^19^ Both scFvs were attached to the respective IgG light chains via a flexible (G_4_S)_3_ linker. The OKT3 scFv was furthermore stabilized with an additional disulfide bridge between the VH and VL domains. The Fc portions of the respective anti-L1CAM IgG contained the L236A, L237A mutations in the hinge region, which have been shown to reduce the effector functions of the Fc part such antibody-dependent cellular cytotoxicity (ADCC), complement-dependent cytotoxicity (CDC) and antibody-dependent cellular phagocytosis (ADCP) by strongly reducing binding to Fcγ receptors and C1q.^20^ As negative control a bispecific antibody with the same IgG (L) scFv format and the same L1CAM binding domain as L1-bsAB1 was constructed, where CD3 binding was abolished by the introduction of targeted mutations.

Surface plasmon resonance experiments showed monovalent binding affinities of 0.15 nM and 0.22 nM towards human L1CAM for L1-bsAB1 and L1-bsAB2, respectively (**Figure 1A**). The association (k_a_) and dissociation (k_d_) rate constants were calculated as k_a_=4.41 × 10^5^ (M-s)^-1^ and k_d_=6.65 × 10^−5^ s^-1^ for L1-bsAB1 and k_a_=3.46 × 10^5^ (M-s)^-1^ and k_d_=7.74 × 10^−5^ s^-1^ for L1-bsAB2 respectively. Both antibodies furthermore showed similar binding to CD3, as shown by flow cytometry using T cells isolated from human PBMC. OKT3 served as a positive control for T cell binding (**Figure 1B**). As expected from the introduced mutations in the hinge region, binding to both isoforms of human FcγRIIIa, the main mediator of ADCC, was markedly reduced when compared to wild-type human IgG1 (**Figure 1A**). Similarly, binding to C1q, the initiator of the CDC pathway, was impaired relative to human IgG1 (**Figure 1A**). FcRn binding on the other hand was not affected by the introduced mutations (**Figure 1A**).

**Figure 1:**
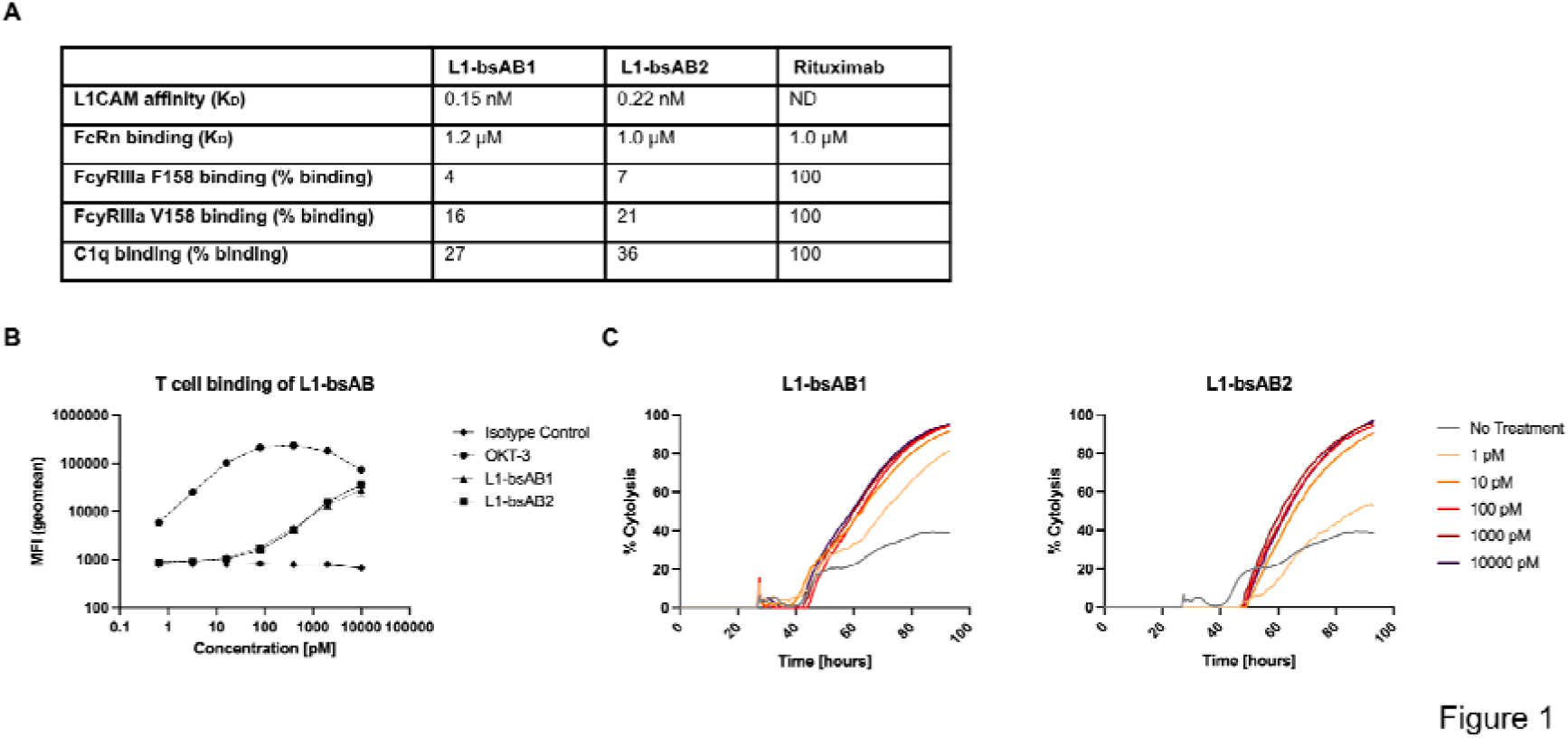
Characterization of L1-bsAB. (**A**) Binding characteristics of L1-bsAB1 and L1-bsAB2 relative to the reference human IgG1 Rituximab were determined by surface plasmon resonance (L1CAM-binding, Fc receptor binding) and ELISA (C1q binding). (**B**) CD3 binding by L1-bsAB1 and L1-bsAB2. T cells isolated from PBMC from healthy donors were incubated with a 7-point concentration series of the bispecific antibodies, OKT3 or human IgG1 isotype control, followed by detection with a PE-conjugated anti-human IgG. Binding of the bispecific antibodies was assessed by quantification of median fluorescent intensity of the PE channel by flow cytometry. Data are presented as mean with standard deviation (SD) from 3 technical replicates using PBMC from a single healthy donor. (**C**) Cytotoxicity of L1-bsAB1 and L1-bsAB2 was determined by impedance measurement of Panc1 cells with a real time cell analysis station. Panc1 cells were seeded 24 h prior to initiation of treatment at indicated concentrations and co-culture with T cells from a single healthy donor at an effector: target ratio of 3:1. Impedance measurement was continued for 72 h after initiation of treatment. Cytotoxicity was calculated from cell indices with the xCelligence Immunotherapy Software. Indicated values represent the mean of two technical replicates.

Next, kinetics of target cell killing were assessed by real-time cell analysis using L1CAM expressing PDAC cell line Panc1 and PBMC from healthy donors as effector cells at an effector: target ratio of 3:1 (**Figure 1C**). Onset of cytotoxic activity was observed 16 h and 24 h after start of treatment with L1-bsAB1 and L1-bsAB2, respectively. For both L1-bsAB, concentrations of 10 pM or higher showed very similar overall target cell killing efficacy, whereas treatment with the lowest concentration of 1 pM showed slower and lower overall cytotoxicity, most notably after treatment with L1-bsAB2.

### L1-bsAB reduce tumor cell viability in 2D PDAC models with effector cells derived from healthy donors

To assess the anti-tumor activity of L1-bsAB1 and L1-bsAB2, L1CAM protein levels of L1- and L1+ variants of the PDAC cell lines Panc1 and Panc89 previously characterized by Brauer et al. were confirmed.^21^ Immunocytochemical staining revealed strong L1CAM positivity in both L1+ variants of Panc1 and Panc89 cells, while both L1- cell variants lacked expression of L1CAM (**Supplemental Figure 1A)**.

To test whether both L1-bsAB mediate target antigen-specific killing, L1- and L1+ variants of both PDAC cell lines were treated with L1-bsAB in the absence or presence of PBMC from healthy donors for 48 h (effector: target ratio of 10:1) (**Figure 2A**). A treatment duration of 48 h was chosen to limit unspecific tumor cell killing by allogeneic effector cells observed in kinetic cytotoxicity experiments (**Figure 1C**). First, we determined the number of remaining live tumor cells (cell-tracker green positive, PI negative) after treatment by automated fluorescence microscopy. While cell viability of L1-PDAC cells was neither impacted by ctrl-bsAB nor L1-bsAB in the presence of PBMC, a significant concentration-dependent reduction of viable L1+ Panc1 and Panc89 cells was determined in co-culture with PBMC after treatment with either L1-bsAB (**Figure 2B**). The strongest effect was observed at 100 pM, the highest concentration of L1-bsAB applied. Treatment reduced viable L1+ PDAC cells to approximately 0.35-0.4-fold for L1-bsAB1 and 0.2-0.5-fold for L1-bsAB2 for Panc1 and Panc89, respectively. L1-bsAB treatment in the absence of PBMC did not impact PDAC cell viability.

**Figure 2:**
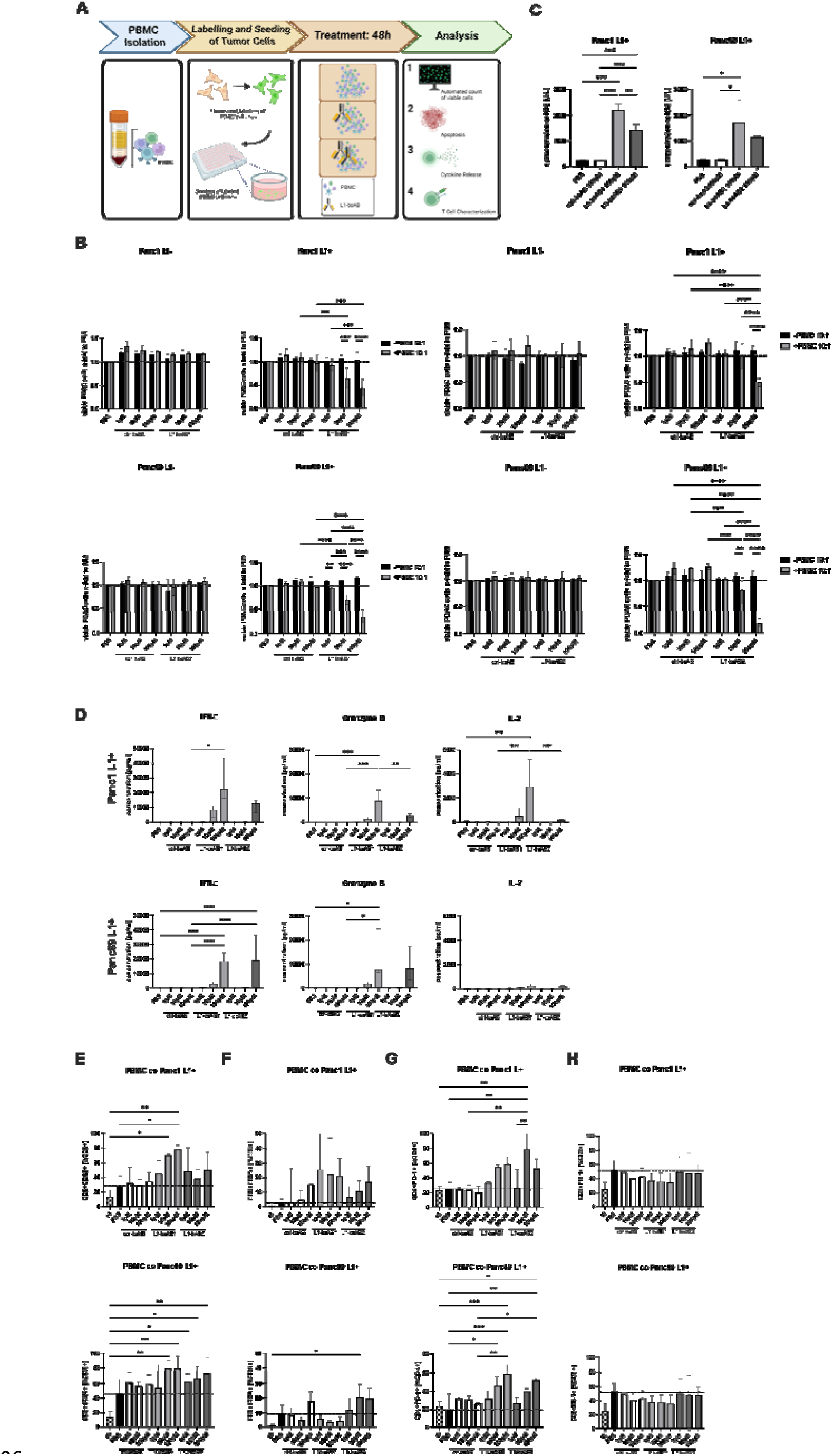
L1-bsAB efficacy in 2D PDAC models with PBMC from healthy donors. (A) Schematic overview of the experimental workflow to determine the number of viable PDAC cells, apoptosis of PDAC cells, release of cytokines by PBMC and expression of activation markers in T cells after treatment with L1-bsAB using PBMC from healthy donors as effector cell population. PDAC cells were stained with cell tracker green to allow for discrimination from PBMC. Stained PDAC cells were seeded and allowed to attach for 24 h. In parallel, PBMC were isolated from healthy donors. Co-culture with PBMC from healthy donors (effector: target ratio 10:1) and treatment with PBS, ctrl-bsAB, L1-bsAB1 or L1-bsAB2 at indicated concentrations were initiated simultaneously and performed for 48 h. Analyses of PDAC cells and PBMC were performed by automated cell imaging, supernatant analyses for ccK18, determination of cytokine release by multiplex assay and flow cytometry of PBMC. (B) Antibody specificity for L1CAM was evaluated by automated cell imaging. PDAC cell lines with differential L1CAM expression (Panc1 L1-, Panc89 L1-, Panc1 L1+, Panc89 L1+) were co-cultured with PBMC from healthy donors (effector: target ratio 10:1) for indicated conditions and treated with PBS, ctrl-bsAB, L1-bsAB1 or L1-bsAB2 for 48 h. Live PDAC cells (Cell tracker green and Hoechst positive, PI negative) were counted with an automated cell imager after treatment. Data is presented as n-fold of viable PDAC cells treated with PBS. (C) Determination of tumor cell specific apoptosis by ccK18 ELISA from culture supernatants. Data is presented as ccK18 level [U/l]. (**D**) Release of IFN-γ, Granzyme B and IL-2 from PBMC co-cultured with Panc1 L1+ (upper panel) and Panc89 L1+ (lower panel) measured by multiplex assay. Data is presented as concentration [pg/ml]. (**E-H**) Flow cytometry analysis of PBMC 48 h after initiation of drug treatment and co-culture with indicated PDAC cells. Staining was performed for T cell activation markers (**E**) CD69, (**F**) CD25 and (**G**+**H**) immunoregulatory receptor PD-1. Data is shown for (**G**) CD4+PD-1 (%CD4) and (**H**) CD8+PD-1+ (%CD8). (**B**) Normality was tested with the Shapiro-Wilk test. A two-way ANOVA followed by Tukey’s multiple comparisons test was applied for normally distributed data. Not-normally distributed data was analyzed by the Kruskal-Wallis test followed by Dunn’s multiple comparisons test. (**C-H**) Normality was tested with the Shapiro-Wilk test. A one-way ANOVA followed by Tukey’s multiple comparisons test was applied for normally distributed data. Not-normally distributed data was analyzed by the Kruskal-Wallis test followed by Dunn’s multiple comparisons test. Data is presented as mean (SD) for normally distributed data and median with interquartile range for not normally distributed data. Every analysis was performed with n=3 independent experiments using PBMC from 3 different donors and significances are indicated by asterisks: * = p<0.05; ** = p<0.01; *** = p<0.001.

To evaluate the efficacy of either L1-bsAB independent of T cell activation kinetics, we additionally assessed L1-bsAB using pre-activated CD8+ T cells as effector cells (**Supplemental Figure 1B**). Here, we already observed significant levels of target cell killing with either L1-bsAB after a treatment duration of 8 h accompanied by more pronounced tumor cell killing for L1-bsAB concentrations of 10 pM and 100 pM.

Altogether, this data underscores that the PDAC cell viability-reducing effect of L1-bsAB is highly specific to the tumor antigen and dependent on the presence of effector cells.

To evaluate whether the reduced PDAC cell viability is related to induction of apoptosis, we determined the levels of caspase-cleaved K18 fragments (ccK18) specific for apoptosis of epithelial/carcinoma cells after 48h co-culture with PBMC from healthy donors and L1-bsAB treatment (**Figure 2C**). In L1-bsAB1 and L1-bsAB2 treated conditions of both L1+ Panc1 and Panc89 cell variants, levels of ccK18 were significantly increased in comparison to control conditions (PBS and ctrl-bsAB), with L1-bsAB1 treatment resulting in overall higher concentrations of ccK18 compared to L1-bsAB2 treatment.

Next, we investigated whether L1-bsAb treatment in the presence of PBMC from healthy donors induces an effector cell response by quantifying the release of cytokines (IFN-γ, IL-2, IL-6, IL-10, IL-17A, IL-6, IL-10) and T cell effector molecules (Granzyme A, Granzyme B, Perforin, Granulysin) into the supernatants by multiplex assay (**Figure 2D, Supplemental Figure 1C**). We observed a dose-dependent release of Granzyme B, IFN-γ and IL-2 after treatment with either L1-bsAB1 or L1-bsAB2, albeit less pronounced for L1-bsAB2, compared to control treatment (**Figure 2D**). Similarly, levels of Granzyme A, Perforin, Granulysin and cytokines IL-6, IL-10 and IL-17A were increased (**Supplemental Figure 1C**). Absolute levels of cytokine release differed between target PDAC cell lines. Overall IL-2 release upon treatment was markedly higher after co-culture with Panc1 than Panc89 cells, while IL-6 was inversely higher upon treatment and co-culture with Panc89 cells (**Figure 2D, Supplemental Figure 1C**).

To determine insights into the kinetics of T cell activation underlying L1-bsAB efficacy, the activation of effector T cells in response to bsAB treatment was assessed by flow-cytometry analysis of activation markers CD69 (early activation) and CD25 (late activation) after 48 h co-culture with L1+ Panc1 or Panc89 cells (**Figure 2E+F**). The gating strategy is depicted in **Supplemental Figure 2**. Overall, CD8+CD69+ and CD8+CD25+ cell populations were increased in control conditions (PBS and ctrl-bsAB) after co-culture with both PDAC cell lines in comparison to pre-treatment conditions. Treatment with either L1-bsAB resulted in a dose-dependent increase of the CD8+CD69+ cell population, with L1-bsAB1 exerting stronger effects (**Figure 2E**). The CD8+CD25+ cell population was also impacted by L1-bsAB treatment demonstrating a more pronounced increase in co-culture with Panc1 than Panc89 cells. (**Figure 2F**).

Since T cell activation may lead to an upregulation of immunoregulatory receptors or ligands limiting T cell response and potentially resulting in T cell exhaustion, we investigated the surface protein levels of PD-1 (**Figure 2G+H**) and PD-L1 (**Supplemental Figure 1D**) in CD4+ and CD8+ T cells after L1-bsAB treatment. While the proportion of PD-1+ cells in CD4+ T cells was not significantly altered by co-culture with either PDAC cell line in the absence of treatment, the proportion of the CD4+PD-1+ cell population increased after treatment with either L1-bsAB (**Figure 2G**), an effect which was not observed for CD8+ T cells (**Figure 2H**). For PD-L1, similar trends were observed as an L1-bsAB-mediated increase in CD4+PD-L1+ cells was determined after co-culture with both PDAC cell lines, while the CD8+PD-L1+ cell population was not relevantly impacted by L1-bsAB treatment (**Supplemental Figure 1D**).

In summary, these data underscore the target antigen and dose-dependent efficacy of both L1-bsAB constructs leading to potent induction of PDAC cell apoptosis along with a pronounced activation and elevated release of T cell effector molecules.

### Determination of L1CAM threshold levels for anti-tumoral effects of L1-bsAB in 2D PDAC models

To determine the required proportion of L1+ tumor cells for a maximum anti-tumoral effect of either L1-bsAB, L1-bsAB treatment was performed with co-cultures of defined ratios of L1+ and L1-PDAC cells with PBMC for 48 h. 24 h after seeding and before starting L1-bsAB treatment, we performed immunocytochemical staining for L1CAM to confirm the defined composition of L1- and L1+ cell variants of Panc1 and Panc89 cells (**Figure 3A**).

**Figure 3:**
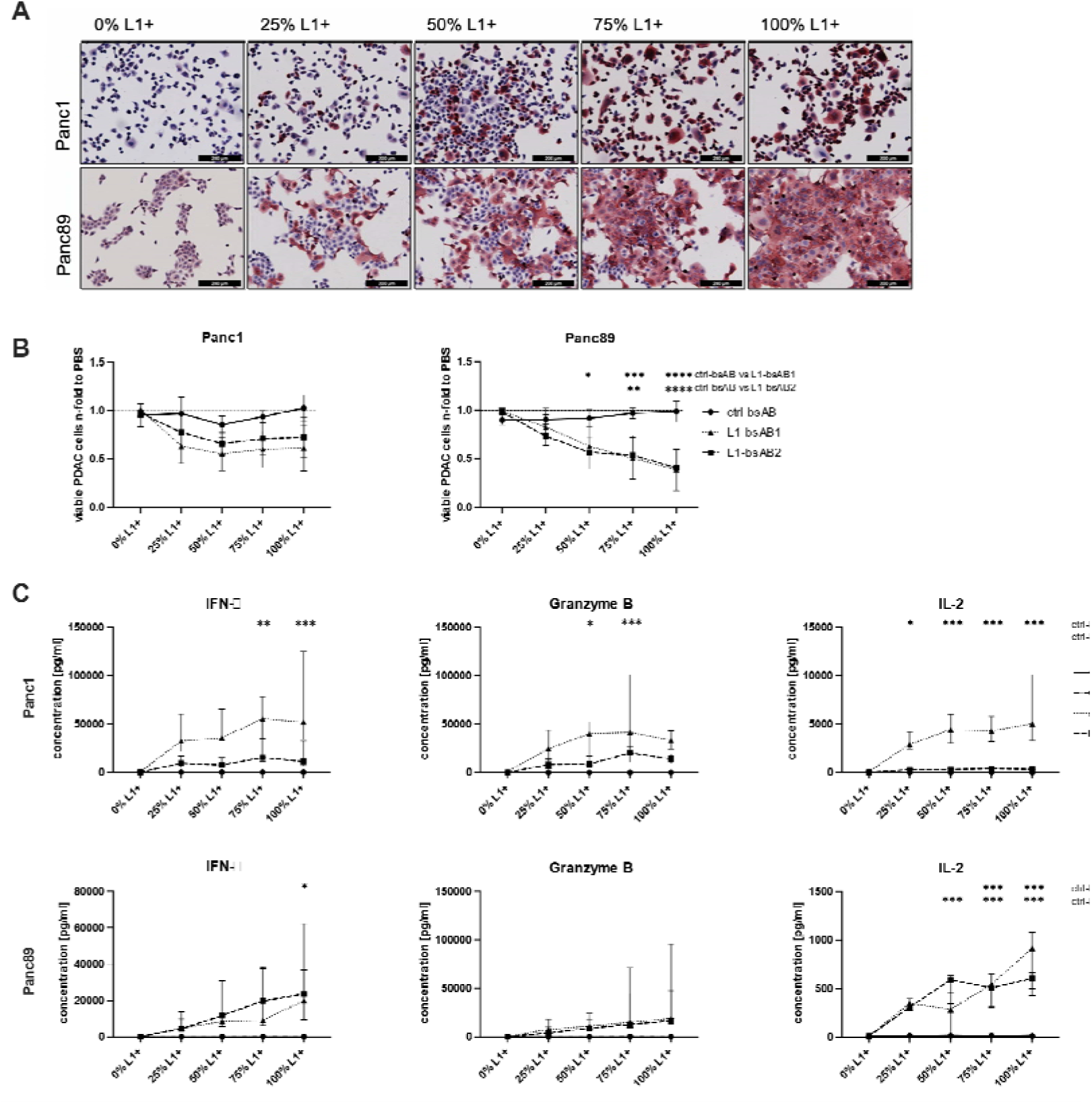
Determining L1CAM threshold levels for L1CAM-bsAB efficacy in 2D PDAC models. (**A-C**) Panc1 and Panc89 were seeded at defined ratios of L1- and L1+ cells. 24 h after seeding, treatment with PBS, ctrl-bsAB, L1-bsAB1 or L1-bsAB2 and co-culture with PBMC from healthy donors was initiated (effector: target ratio 10:1). All antibodies were applied at a concentration of 100 pM. (**A**) Representative images of immunocytochemical staining of L1CAM (red) of PDAC cells seeded at indicated ratios of L1- and L1+ on coverslips. Staining was performed 24 h after seeding. Nuclei were stained with Mayer’s hemalum (blue). Scale bar: 200 µm. (**B**) Determination of viable PDAC cells 48 h after treatment and initiation of co-culture. Cells were stained with Hoechst (nuclei) and PI (dead cells). Live PDAC cells (Cell tracker green and Hoechst positive, PI negative) were counted using an automated cell imager. Data is presented as n-fold of cells treated with PBS (indicated by dashed line). Data from n=4 independent experiments using PBMC from 4 different donors. (**C**) Release of IFN-γ, Granzyme B and IL-2 from PBMC co-cultured with Panc1 (upper panel) and Panc89 (lower panel) and treated with indicated drugs measured by multiplex assay. Data is presented as concentration [pg/ml]. Data from n=3 independent experiments using PBMC from 3 different donors. (**B+C**) Normality was tested with the Shapiro-Wilk test. A one-way ANOVA followed by Tukey’s multiple comparisons test was applied for normally distributed data. Not-normally distributed data was analyzed by the Kruskal-Wallis test followed by Dunn’s multiple comparisons test. Data is presented as mean (SD) for normally distributed data and median with interquartile range for not normally distributed data. Significances are indicated by asterisks: * = p<0.05; ** = p<0.01; *** = p<0.001.

We then assessed the efficacy of L1-bsAB1 and L1-bsAB2, each applied at a concentration of 100 pM and an effector to target ratio of 10:1 in the different PDAC cell populations with PBMC. Drug effects on the number of viable cells differed between target cell lines Panc1 and Panc89 (**Figure 3B**). In Panc1 cells, we already observed a trend towards a reduction of tumor cells to approximately 0.6-fold with L1-bsAB1 and 0.8-fold with L1-bsAB2 compared to PBS control, when 25% of target cells were L1+ at the timepoint of seeding. Of note, in Panc1 cells, further increasing the number of target antigen-expressing cells did not result in relevant improvements in drug efficacy. Contrastingly, in Panc89 cells the number of viable cells continuously decreased with increasing proportions of L1+ cells. The proportion of viable cells ranged from approximately 0.8-fold and 0.7-fold with 25% L1+ cells after treatment with L1-bsAB1 and L1-bsAB2, respectively, and 0.4-fold for both L1-bsAB with a proportion of 100% L1+ cells when compared to PBS control condition.

Overall, this data further confirms that both L1-bsAB are more efficient in Panc89 than Panc1 cells and that the efficacy of either L1-bsAB is dependent on the number of target antigen-expressing tumor cells. The observation that in Panc1 cells the maximum antibody efficacy was already observed when 25% of the tumor cell population exhibits the target antigen may pinpoint to potential resistance mechanisms in this tumor cell line.

To elucidate the effects of L1-bsAB treatment on T cell activity depending on the number of L1+ target cells, we determined levels of secreted T cell effector molecules and cytokines. Despite the plateau of the reduced cell viability observed in Panc1 cells with 25% of L1+ target cells, we observed a L1+ cell number dependent increase of the assessed effector molecules and cytokines in supernatants of co-cultures with both PDAC cell lines **(Figure 3C, Supplemental Figure 3**). Moreover, along with the comparable anti-tumor cell effects of both L1-bsAB (**Figure 3B**) the release of T cell effector molecules (Granzyme A, Granzyme B, Perforin, Granulysin) was similarly elevated, while the cytokine release was also elevated, but lower upon L1-bsAB2 treatment (IFN-γ, IL-2, IL-10, IL-17A). Taken together, L1-bsAB1 and L1-bsAB2 both induce relevant reduction of viable PDAC cells already at relatively low proportion of L1+ target cells of 25% with the release of T cell effector molecules and cytokines being dependent on the L1-bsAB construct.

### L1-bsAB reduce tumor cell viability in 2D PDAC models with patient-derived effector cells

Tumor-infiltrating T cells and T cells circulating in the peripheral blood of PDAC patients express increased levels of immunoregulatory receptors including PD-1 compared to healthy donors and may therefore be functionally impaired.^22^ Thus, we next assessed L1-bsAB efficacy in co-cultures of L1+ PDAC cells and PDAC patient-derived PBMC as effector population. Patient characteristics are described in detail in **Table 1**.

**Table 1:**
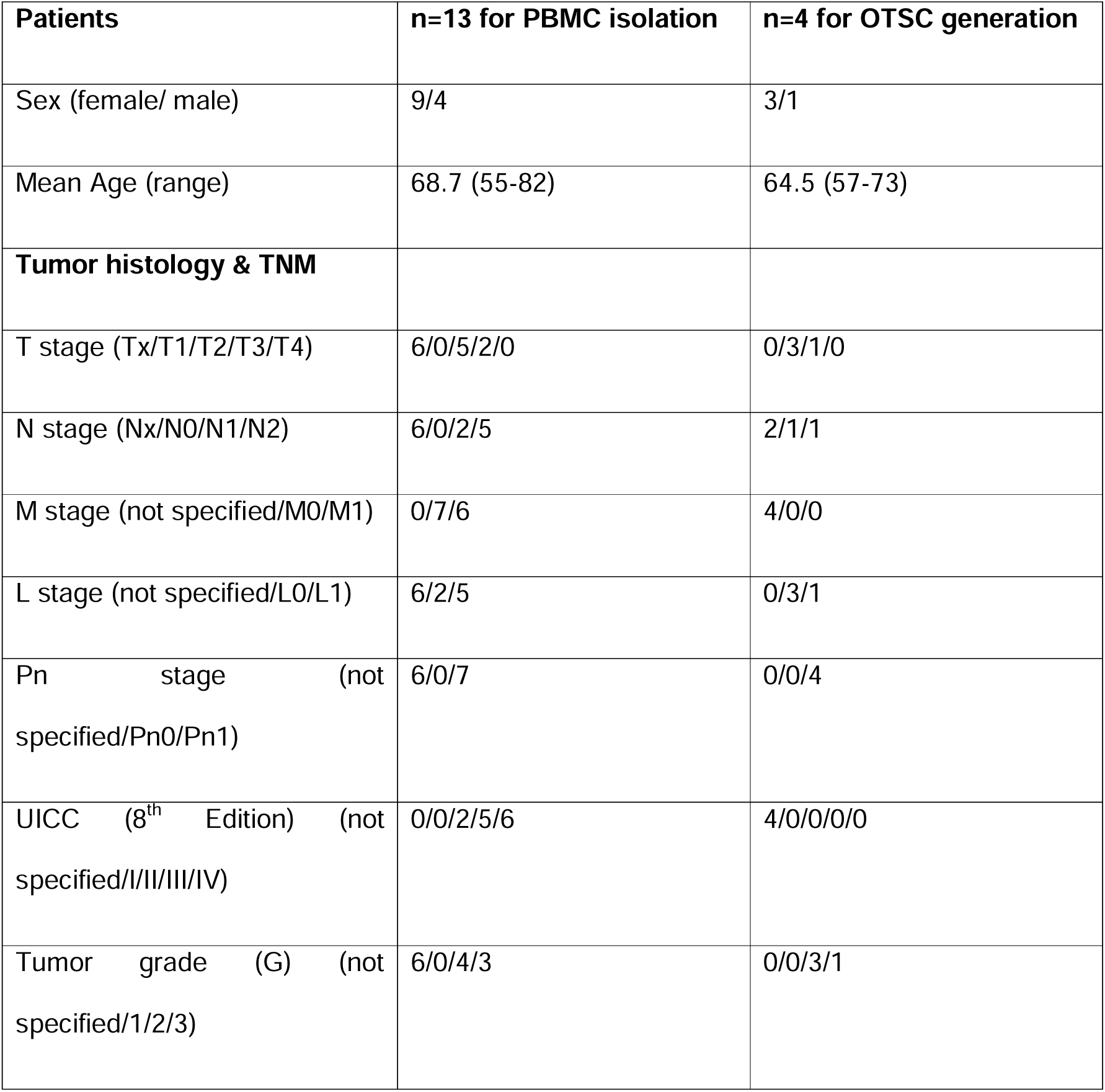
Clinical and pathological characteristics of PDAC patients whose PBMC were used for co-culture experiments or whose tumor tissue was used for generation of OTSC. The Classification of Malignant Tumors according to the Union Internationale Contre le Cancer (UICC) was determined based on the pathological tumor stage (T), regional lymph nodal stage (N), distant metastatic stage (M) as well as invasion into lymphatic vessels (L) and adjunct nerves (Pn). Tumor grade (G) describes the cellular differentiation.

| **Patients** | **n=13 for PBMC isolation** | **n=4 for OTSC generation** |
| --- | --- | --- |
| Sex (female/ male) | 9/4 | 3/1 |
| Mean Age (range) | 68.7 (55-82) | 64.5 (57-73) |
| **Tumor histology & TNM** |  |  |
| T stage (Tx/T1/T2/T3/T4) | 6/0/5/2/0 | 0/3/1/0 |
| N stage (Nx/N0/N1/N2) | 6/0/2/5 | 2/1/1 |
| M stage (not specified/M0/M1) | 0/7/6 | 4/0/0 |
| L stage (not specified/L0/L1) | 6/2/5 | 0/3/1 |
| Pn stage (not specified/Pn0/Pn1) | 6/0/7 | 0/0/4 |
| UICC (8<sup>th</sup> Edition) (not specified/I/II/III/IV) | 0/0/2/5/6 | 4/0/0/0/0 |
| Tumor grade (G) (not specified/1/2/3) | 6/0/4/3 | 0/0/3/1 |

As before, cell viability was determined 48 h after initiation of co-culture at an effector to target ratio of 10:1 and treatment with either ctrl-bsAB, L1-bsAB1 or L1-bsAB2 at the indicated concentrations. We observed an overall higher variability of anti-tumorigenic efficacy of either L1-bsAB with PBMC from PDAC patients compared to PBMC from healthy donors (**Figure 2B and 4A**). Moreover, higher concentrations of either L1-bsAB had to be applied to significantly reduce the number of viable PDAC cells with concentrations up to 10 nM of L1-bsAB1 or L1-bsAB2 (**Figure 2B and 4A**). In line with the observations made using PBMC from healthy donors, cell viability of Panc89 cells was already noticeably reduced at 100 pM compared to Panc1 cells.

**Figure 4:**
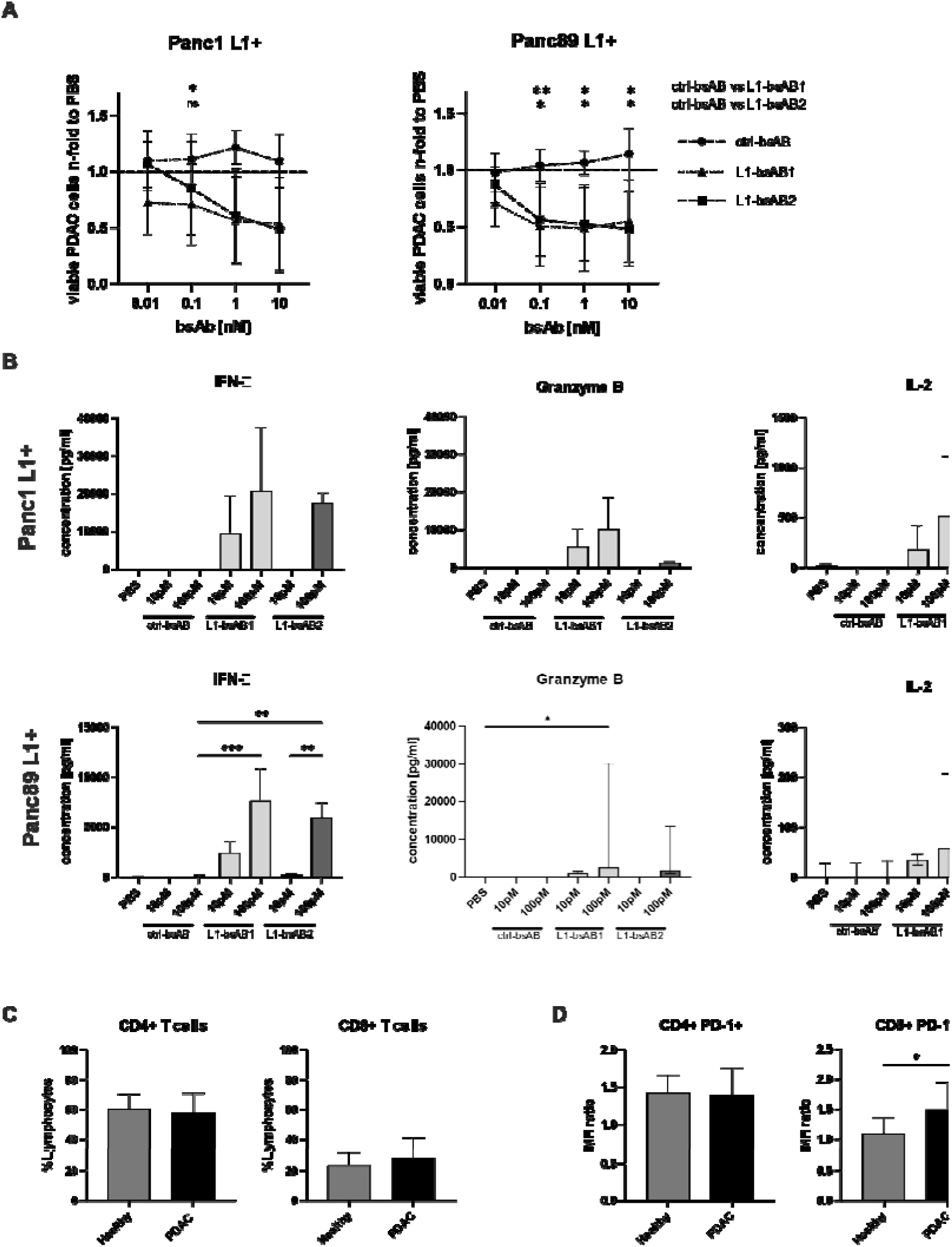
L1-bsAB efficacy in 2D PDAC models using PBMC from PDAC patients as effector population. (**A**) Determination of viable PDAC cells 48 h after treatment with ctrl-bsAB, L1-bsAB1 or L1-bsAB2 at indicated concentrations and initiation of co-culture with PBMC from PDAC patients (effector: target ratio 10:1). Live PDAC cells (Cell tracker green and Hoechst positive, PI negative) were counted using an automated cell imager. Data is presented as n-fold of cells treated with PBS (indicated by dashed line). Data from n=5 independent experiments using PBMC from 5 different donors. (**B**) Release of IFN-γ, Granzyme B and IL-2 from PBMC of PDAC patients co-cultured with Panc1 L1+ (upper panel) and Panc89 L1+ (lower panel) and treated with indicated drugs measured by multiplex assay. Data is presented as concentration [pg/ml]. Data from n=3 independent experiments using PBMC from 3 different donors. (**C+D**) Flow cytometry analysis of PBMC from healthy donors (healthy) (n=10) and PDAC patients (PDAC) (n=8). (**C**) Staining was performed for expression of CD4 and CD8. Data is shown as percentage of positive cells in total lymphocytes (%Lymphocytes). (**D**) Staining for the immunoregulatory receptor PD-1 was performed and its expression was determined for the respective T cell population (CD4+ or CD8+). Data is shown as MFI-Ratio normalizing the intensity of specific immunofluorescence staining to the staining intensity of the respective isotype control antibody and calculating the ratio of median fluorescence intensities. (**A-B**) Normality was tested with the Shapiro-Wilk test. A one-way ANOVA followed by Tukey’s multiple comparisons test was applied for normally distributed data. Not-normally distributed data was analyzed by the Kruskal-Wallis test followed by Dunn’s multiple comparisons test. (**C-D**) Normality was tested with the Shapiro-Wilk test. Data was analyzed by the Welch’s t-test. (**A-D**) Data is presented as mean (SD) for normally distributed data and median with interquartile range for not normally distributed data. Significances are indicated by asterisks: * = p<0.05; ** = p<0.01; *** = p<0.001.

In accordance with the reduced cell viability after L1-bsAB treatment in co-cultures of PDAC cells and PDAC patient-derived PMBC, we observed a dose-dependently increasing release of T cell effector molecules and cytokines in cell culture supernatants (**Figure 4B, Supplemental Figure 4**). Furthermore, the lower anti-tumorigenic efficacy of L1-bsAB1 and L1-bsAB2 in the presence of PDAC patient-derived PBMC was accompanied by a lower concentration of T cell effector molecules and cytokines compared to co-culture with PBMC derived from healthy donors (**Figure 2D and 4B**). Notably, the concentrations of IFN-γ, IL-2 (**Figure 2D and 4B**), Granzyme A and IL-17A (**Supplemental Figure 1C and 4**) were lower in supernatants from co-culture with PDAC patient-derived PBMC compared to co-culture with PBMC from healthy donors. Contrastingly, levels of T cell effector molecules Granzyme B, Perforin and Granulysin were only slightly reduced or reached comparable levels (**Figure 4B, Supplemental Figure 4**).

Finally, we compared composition and PD-1 status of CD4+ and CD8+ T cells of healthy donors and PDAC patient-derived PBMC. While the proportion of CD4+ and CD8+ T cell populations was comparable between healthy donors and PDAC patients (**Figure 4C**), CD8+ but not CD4+ positive T cells exhibited significantly higher surface protein levels of PD-1 (**Figure 4D**) in line with previously published data.^22^ Overall, the data demonstrate potent anti-tumor effects of L1-bsAB with PDAC patient-derived effector cells albeit with a higher variability than observed with PBMC from healthy donors.

### L1-bsAB reduce tumor cell viability in PMF-enriched 3D PDAC cell spheroids

Besides the 3D context, PDAC cell biology is massively shaped by its TME. Especially, PMF and macrophages, being two prominent TME cell populations, contribute to the immunosuppressive milieu, thereby inhibiting T cell effector function.^23^ To explore whether L1-bsAB is effective not only in the 3D context but also in the presence of those immunosuppressive stromal cells, a 3D Panc89 cell spheroid model enriched with different stromal cell populations was employed. To mimic the heterogeneous L1CAM expression observed in patient tumors, Panc89 parental cells comprehensively characterized by Brauer et al. were used for further experiments.^21^

In the first model, Panc89 parental cells were either seeded alone (mono-cultured) or together with PMF at two different ratios (PDAC cells: PMF of 1:1 (co1) or 1:2 (co2)) in ultra-low attachment plates and allowed to form 3D spheroids for 24h. Respective mono-culture spheroids contained the identical number of tumor cells only (mono1 same number as co1 and mono2 same number as co2).

After successful formation of spheroids, PBMC from healthy donors were added at a ratio of 10:1 (PBMC: non-effector cells) and treatment with L1-bsAB1 or L1-bsAB2 was initiated at different concentrations for 48 h (**Figure 5A**).

**Figure 5:**
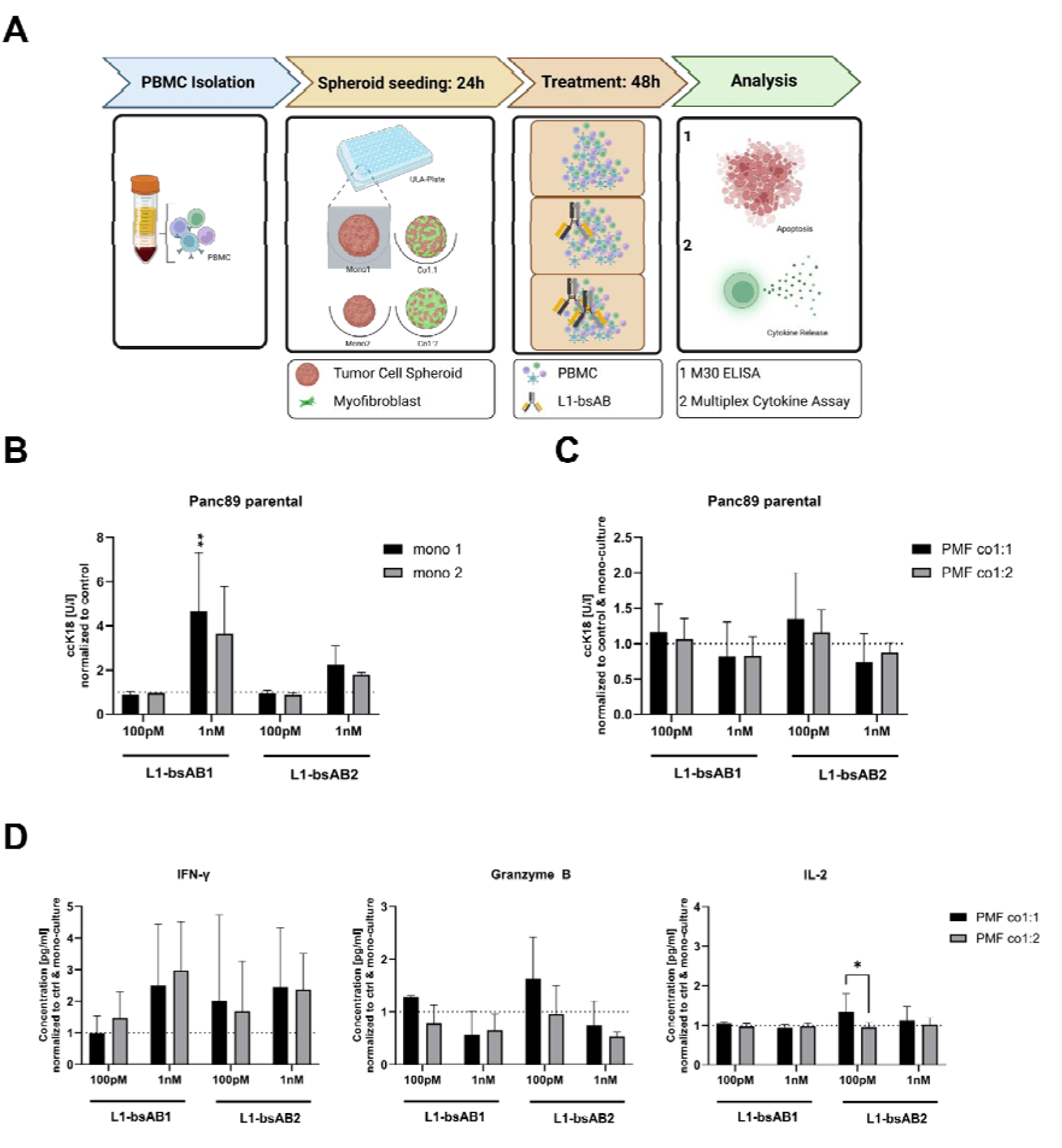
L1-bsAB efficacy in PMF-enriched 3D PDAC cell spheroids. (A) Schematic overview of the experimental workflow to determine the impact of PMF on L1-bsAB treatment in a 3D PDAC spheroid co-culture model. Panc89 parental cells were seeded alone (mono) or in co-culture with different amounts of PMF (Panc89: PMF ratio 1:1 or 1:2) to form spheroids. Mono1 and mono2 represent mono spheroids with the same number of tumor cells used for co1 and co2. After L1-bsAB effects were determined by quantification of tumor cell specific apoptosis via ccK18 level, release of cytokines by PBMC from healthy donors was analyzed via multiplex analysis. **(B-C)** Determination of tumor cell specific apoptosis by ccK18 ELISA from culture supernatants in (**B**) mono-cultured and (**C**) co-cultured spheroids. (**D**) Release of IFN-γ, Granzyme B and IL-2 from PBMC from healthy donors co-cultured with Panc89/PMF co-culture spheroids and treated with indicated drugs measured by multiplex assay. Data is presented as concentration [pg/ml] normalized to ctrl-bsAB treatment and mono-culture spheroids. (**B-D**) Normality was tested with the Shapiro-Wilk test. A two-way ANOVA followed by Tukey’s multiple comparisons test was applied for normally distributed data. Not-normally distributed data was analyzed by the Kruskal-Wallis test followed by Dunn’s multiple comparisons test. Every analysis was performed with n=3 independent experiments using PBMC from 3 different donors. Data is presented as mean with SD for normally distributed data and median with interquartile range for not normally distributed data. Significances are indicated by asterisks: * = p<0.05; ** = p<0.01.

First, tumor cell apoptosis was determined by detecting ccK18 level (**Figure 5B+C**). L1-bsAB1 significantly induced apoptosis, especially in mono spheroids, indicated by elevated ccK18 level (approximately 4-fold to control conditions), in mono spheroids at higher concentration, whereas apoptosis was only slightly induced after treatment with L1-bsAB2 (**Figure 5B**). Compared to 2D conditions, higher concentrations of L1-bsAB1 were required in 3D PDAC models to significantly induce apoptosis in PDAC cells. irrespective of the presence of PMF.

Of note, co-culture with PMF did not impair treatment response to both L1-bsAB compared to mono-cultured spheroids indicated by the almost unchanged normalized ccK18 levels of PMF enriched co-cultures compared to mono-cultured spheroids (**Figure 5C**).

We next investigated the levels of secreted T cell effector molecules and cytokines. As observed in 2D cultures, L1-bsAB treatment led to an elevated release of IFN-γ and Granzyme B in supernatants of mono-cultured Panc89 cell spheroids, with L1-bsAB1 exerting stronger effects compared to L1-bsAB2. In contrast, IL-2 was hardly affected by L1-bsAB treatment under 3D conditions (**Supplemental Figure 5A)**.

IFN-γ secretion was further increased in supernatants from PMF-enriched 3D PDAC spheroids upon treatment with either L1-bsAB compared to control treatment and mono-cultured spheroids (**Figure 5D**). In contrast, Granzyme B levels were only slightly increased after L1-bsAB treatment at 100 pM and in PMF co1:1 condition, while at a L1-bsAB concentration of 1 nM lower amounts of Granzyme B were observed under both PMF enriched conditions compared to control and mono-cultured spheroids. Finally, PMF did almost not impact L1-bsAB mediated release of IL-2 (**Figure 5D**).

In contrast to 2D cultures, only slight changes in IL-2 levels could be observed in 3D cultures without PMF (**Supplemental Figure 5A**) and also PMF did not impact IL-2 level much, even though PMF co-culture 1:1 revealed an increase in IL-2 after 100 pM treatment of L1-bAB2, which might be caused by the reduction in the corresponding mono-culture (**Figure 5D and Supplemental Figure 5A**).

Interestingly, 3D co-culture reduced secretion of IL-6 and IL-10 after L1-bsAB1 treatment, while secretion of Granzyme A was slightly increased. Furthermore, secretion of Perforin, Granulysin, and IL-17A was not affected by co-culture (**Supplemental Figure 5C**).

Overall, these data demonstrate the efficacy of L1-bsAB in 3D PDAC cell spheroid models and show that PMF do not impair the efficacy of this TCE under the applied co-culture conditions.

### L1-bsAB also reduce tumor cell viability in M1- or M2- macrophage-enriched 3D PDAC cell spheroids

Next, we assessed the influence of different macrophage cell populations on the efficacy of L1-bsAB treatment. For this purpose, we set up 3D co-culture spheroids as described above with Panc89 parental cells exhibiting heterogeneous L1CAM expression and M1-like or M2- like macrophages at two different ratios. The latter cells were generated through stimulation of CD14+ cells, isolated from peripheral blood of healthy donors, with GM-CSF or M-CSF, respectively (**Figure 6A**).

**Figure 6:**
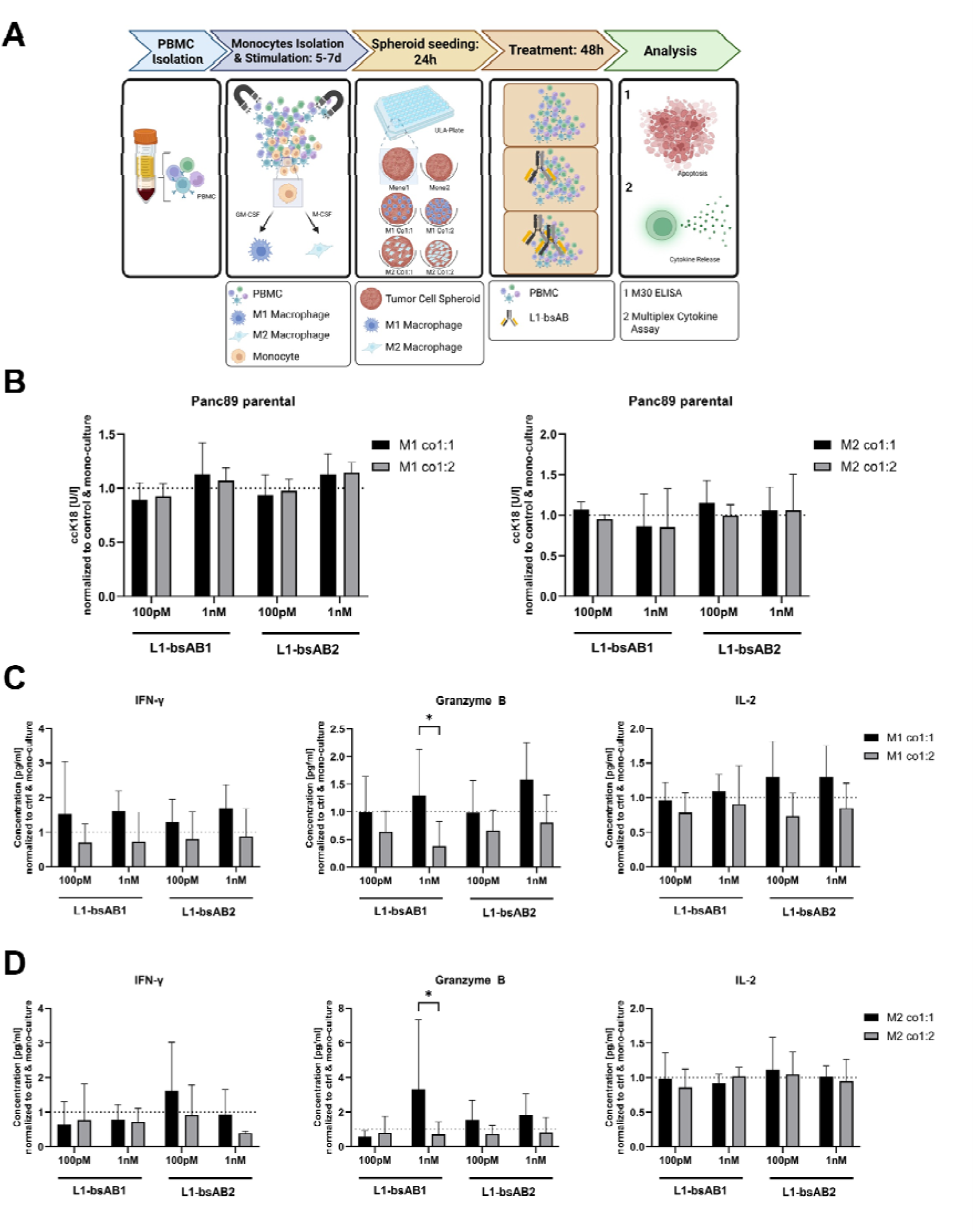
L1-bsAB efficacy in macrophage enriched 3D PDAC cell spheroids. (**A**) Schematic overview of the experimental workflow to determine the impact of M1-like and M2- like macrophages on L1-bsAB treatment in a 3D PDAC spheroid co-culture model. Panc89 parental cells were seeded alone (mono) or in co-culture with different amounts of macrophages (Panc89: macrophage ratio 1:1 or 1:2) to form spheroids. After treatment and co-culture with PBMC for 48h, L1-bsAB efficacy was determined by analysis of tumor cell specific apoptosis via ccK18 level and release of cytokines by PBMC from healthy donors was analyzed via multiplex analysis. **(B)** Determination of tumor cell specific apoptosis by ccK18 ELISA of supernatants from co-cultured and treated spheroids. Release of IFN-γ, Granzyme B and IL-2 from PBMC from healthy donors co-cultured with Panc89 spheroids enriched with **(C)** M1-like macrophages or **(D)** M2- like macrophages and treated with indicated drugs measured by multiplex assay. Data is presented as concentration [pg/ml] normalized to ctrl-bsAB treatment and mono-cultured spheroids. (**B-D**) Normality was tested with the Shapiro-Wilk test. A two-way ANOVA followed by Tukey’s multiple comparisons test was applied for normally distributed data. Not-normally distributed data was analyzed by the Kruskal-Wallis test followed by Dunn’s multiple comparisons test. Every analysis was performed with n=4 independent experiments using PBMC and monocytes from 4 different donors. Data is presented as mean (SD) for normally distributed data and median with interquartile range for not normally distributed data. Significances are indicated by asterisks: * = p<0.05.

Although macrophages did hardly impact L1-bsAB mediated PDAC cell apoptosis (**Figure 6B**), the presence of the higher ratios of M1-like macrophages (1:2) negatively regulated levels of T cell effector molecules including secreted IFN-γ, Granzyme B and IL-2 upon L1-bsAB treatment (**Figure 6C**) underpinning the immunomodulatory effects of macrophages. Similar effects were observed for IFN-γ and Granzyme B in 3D PDAC cell spheroids enriched with M2- like macrophages (**Figure 6D**). Also, secretion of IL-6, IL-10 and Granzyme A was reduced after co-culture especially with M2- like macrophages and treatment with L1-bsAB1 (**Supplemental Figure 6A+B**).

Altogether this data indicates that macrophages, even of an anti-inflammatory M2- like phenotype, do not impair the anti-PDAC cell efficacy of L1-bsAB.

### L1-bsAB shows activity in PDAC organotypic slice cultures with maintained PDAC tumor microenvironment

Finally, to assess L1-bsAB efficacy in a model with structurally preserved tissue architecture andTME, we used patient-derived PDAC OTSCs (**Figure 7**). OTSCs were prepared from freshly resected PDAC specimens (n=4, treatment-naïve). Patient characteristics, including histopathological staging, are described in **Table 1**.

**Figure 7:**
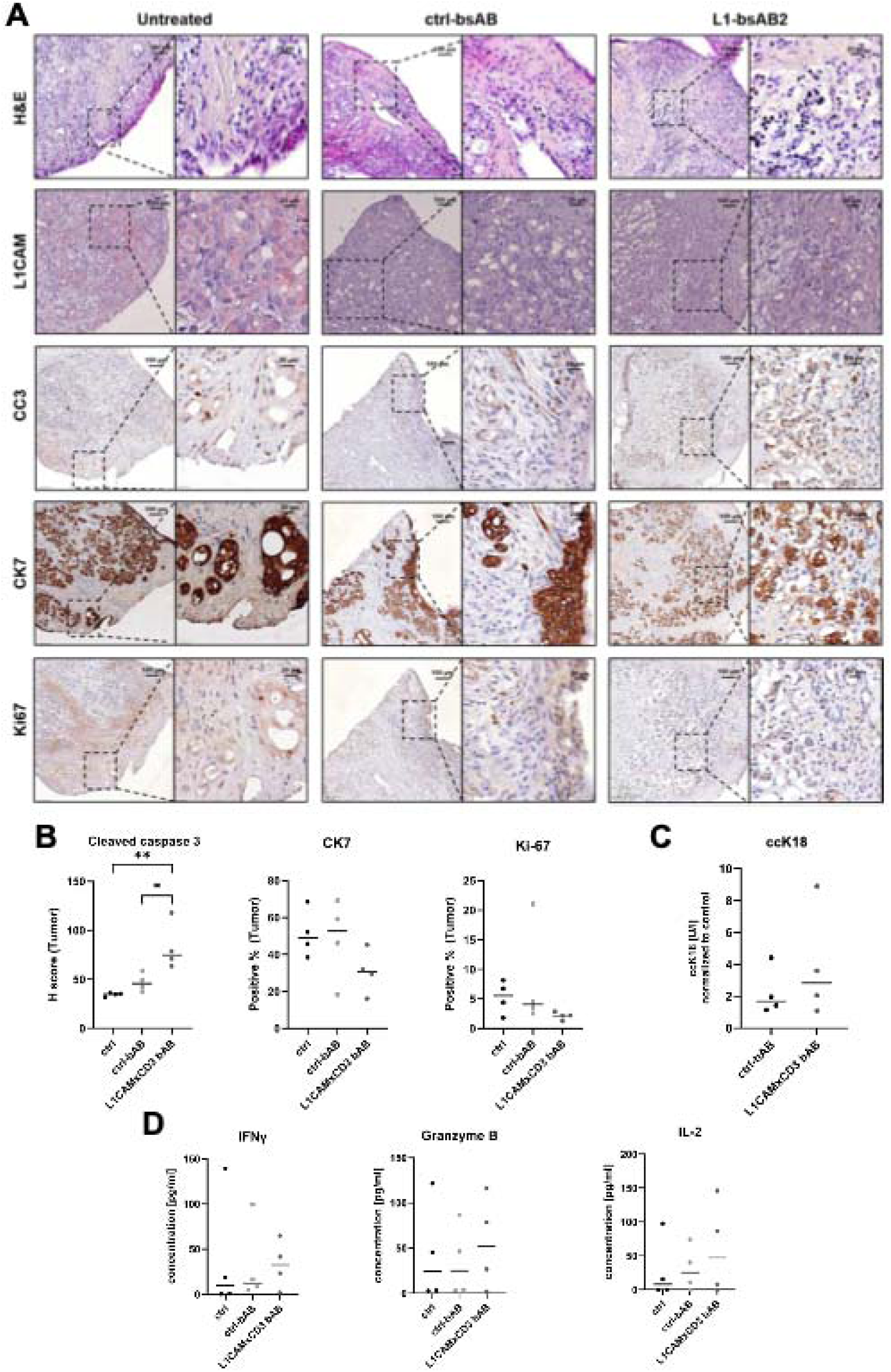
Efficacy of L1-bsAB2 efficacy in OTSCs. **(A**) Representative images of H&E and immunohistochemical staining (L1CAM, cleaved caspase 3 (CC3), cytokeratin 7 (CK7) and Ki-67) of OTSC from one patient. OTSC were treated with PBS (ctrl), ctrl-bsAB or L1-bsAB2 for 48 h. **(B)** L1-bsAB efficacy was assessed by analysis of CC3, CK7, Ki-67 and (**C**) tumor cell specific apoptosis via determination of ccK18 level **(D)** Release of IFN-γ, Granzyme B and IL-2 after L1-bsAB2 treatment into the culture supernatant was measured by multiplex assay. (**B-D**) Data is presented as mean for normally distributed data and median for not normally distributed data with data points representing individual OTSCs. Normality was tested with the Shapiro-Wilk test. A one-way ANOVA followed by Tukey’s multiple comparisons test was applied for normally distributed data. Not-normally distributed data was analyzed by the Kruskal-Wallis test followed by Dunn’s multiple comparisons test. Every analysis was performed with n=4 independent experiments using OTSC from 4 different PDAC patients. Significances are indicated by asterisks: * = p<0.05; ** = p<0.01. Scalebar 100 μm (right) and 20 μm (left).

First, L1CAM expression in PDAC OTSC was determined. In all untreated OTSC samples, L1CAM-positive tumor cells could be detected. Overall, L1CAM protein levels were assessed as comparably low (**Figure 7A**). To evaluate anti-PDAC cell effects of L1-bsAB2 treatment in comparison to control conditions (untreated control = ctrl, ctrl-bsAB treated), we determined the tumor cell content (H&E and Cytokeratin (CK) 7), calculated the proportion of apoptotic tumor cells (cleaved caspase (CC) 3^+^) and the proportion of proliferating tumor cells (Ki67^+^) based on immunohistological quantification (**Figure 7A+B**) and analyzed epithelial/carcinoma cell apoptosis via detection of ccK18 levels in culture supernatants (**Figure 7C**). We observed a trend towards higher apoptosis induction in L1-bsAB2 treated OTSCs indicated by moderately increased ccK18 levels in the supernatants compared to control (**Figure 7C**). In line, CC3 protein levels were significantly higher in L1-bsAB2 treated OTSC compared to control OTSCs, supporting the ability of L1-bsAB2 to induce PDAC cell apoptosis under these TME conditions (**Figure 7B**). In parallel, we observed a trend towards decreased expression of CK7 and less Ki67^+^ cells after treatment with L1-bsAB2, further underlining reduction of total tumor cell content along with a decrease of tumor cell proliferation (**Figure 7B**). Additionally, multiplex analysis of OTSC supernatants revealed a trend towards increased levels of IFN-γ, Granzyme B, IL-2 and IL-6 indicating T cell activation by L1-bsAB2 treatment under these TME conditions (**Figure 7D, Supplemental Figure 7**).

Overall, L1-bsAB2 treatment showed tangible anti-tumor efficacy accompanied by T cell activation in OTSCs despite low L1CAM status of tumor cells present in the investigated OTSCs.

## Discussion

L1CAM expression, mostly located at the tumor invasive front, has been established as a negative prognostic marker for PDAC.^24^ As a high proportion of primary PDAC tissues (approximately 80%) and liver metastases express L1CAM and L1CAM mediates multiple pro-tumorigenic processes in PDAC, it has emerged as a promising target molecule for immunotherapeutic strategies in PDAC.^9^ In this study, we developed two novel T cell engaging L1-bsAB and investigated both in diverse 2D and 3D PDAC models considering the altered immune state of PDAC patients as well as different aspects of the complex TME. In contrast to a previously published L1CAMxCD3 TCE targeting the L1CAM epitope CE7 with a chimeric L1CAM-binding arm ^25^, L1-bsAB1 and L1-bsAB2 are fully humanized and bind to alternative L1CAM epitopes AFF4 and AFF1, respectively. L1-bsAB2 also relies on an alternative CD3 binding domain (humanized SP34). Despite these differences in their design, characterization of both L1-bsAB revealed very similar properties regarding L1CAM affinity, binding to FcRn, FcγRIIa F158, FcγRIIa V158, C1q and T cell binding. Contrastingly, we observed differences in the kinetics of target cell killing. Here, L1-bsAB1 treatment resulted in faster target cell killing than treatment with L1-bsAB2. This observation highly likely reflects subtle differences in T cell activation kinetics mediated by the binding to different CD3 epitopes.

Next, efficacy and target specificity of L1-bsAB were assessed in well-characterized variants of PDAC cell lines Panc1 and Panc89 expressing differential levels of L1CAM (L1- or L1+) using PBMC from healthy donors as effector cell population. High L1CAM-specificity of L1-bsAB1 and L1-bsAB2 was confirmed, as the number of viable target cells was significantly reduced when applied to L1CAM expressing cell variants (Panc1 L1+ and Panc89 L1+) while no effect of L1-bsAB treatment on cell viability was detectable in Panc1 L1- and Panc89 L1- cell variants. The ctrl-bsAB characterized by high L1CAM affinity but lacking a functional CD3-binding domain did not exert relevant anti-tumorigenic effects under these experimental conditions. Additionally, neither L1-bsAB reduced tumor cell viability in the absence of effector cells, both underscoring the necessity of T cell activity via CD3 for L1-bsAB anti-tumor efficacy at the indicated range of concentrations.

Having established L1CAM specificity of both L1-bsAB, we focused on preclinical PDAC models with PDAC cells exhibiting high L1CAM expression (L1+) for further evaluation in 2D models. Using PBMC from healthy donors or PDAC patients as effector cells, we could show that apoptosis was induced in target cells upon L1-bsAB treatment representing a relevant mechanism of cell death. Additionally, we observed dose-dependent activation of T cells by L1-bsAB treatment. As expected due to its slower T cell activation kinetics, L1-bsAB2 had to be applied at higher doses to achieve similar overall levels of tumor cell killing 48h after treatment. Overall, results with PBMC from PDAC patients showed a higher variability and a trend towards reduced efficacy with both L1-bsAB. This may be explained by a higher level of exhausted T cells characterized by increased expression of immune checkpoint proteins, not only locally within the tumor but also circulating systemically. Previous studies demonstrated altered immunophenotypes in circulating T cells in PDAC patients displaying increased expression of immune checkpoint proteins and a shift towards a higher proportion of regulatory T cell populations.^22,26^ In line with these findings, our study revealed significantly elevated PD-1 levels in CD8+ T cells from PDAC patients compared to those from healthy donors. T cell activation upon L1-bsAB treatment was accompanied by a dose-dependent release of T cell effector molecules and cytokines from PBMC of healthy donors as well as PDAC patients. For most investigated molecules, a trend towards lower release from patient-derived PBMC emerged, again pointing towards a functionally impaired T cell population in PDAC patients. Overall, our data underscores the added value of using patient-derived effector cells for TCE efficacy testing to consider the altered functional phenotype of effector cell populations in cancer patients.

Co-culture experiments with defined percentages of L1- and L1+ PDAC cells revealed a target cell specific response to L1-bsAB. In Panc1 cells, target cell killing reached a plateau for both L1-bsAB, when the initially seeded L1+ population exceeded 25% of cells.

Contrastingly, in Panc89 cells treatment response consistently increased with rising percentage of L1+ cells. This finding highlights the potential relevance of target cell intrinsic factors as modifiers of TCE efficacy. Therefore, definition of universal threshold values defined by a percentage of antigen positive target cells may prove difficult as they do not account for individual tumor characteristics modifying response to bsAB. However, future clinical investigation should further assess potential correlations between L1CAM expression and therapy efficacy to determine, if a respective threshold level can be established as a predictive marker for L1-bsAB therapy.

In co-cultures with both PDAC cell lines, the release of T cell effector molecules and cytokines into the culture supernatant depended on the percentage of L1+ target cells. Despite similar anti-tumor efficacy of L1-bsAB1 and L1-bsAB2, we detected higher concentrations of multiple cytokines in the supernatant of cells treated with L1-bsAB1. This finding reflects the differences in the design of both L1-bsAB resulting in differences in effector cell activation kinetics and potentially overall T cell activation levels. Achieving optimal levels of cytokine release is important for amplification of local immune activation, while suboptimal high levels may counterproductively result in local T cell exhaustion or systemic cytokine release syndrome (CRS).^27^ To this end, published data indicate that a bsAB design characterized by high affinity for the selected tumor-antigen and moderate to low CD3 affinity may sustain potent anti-tumor activity, while mitigating cytokine release.^28,29^

To further evaluate the efficacy of L1-bsAB in PDAC cell models that mimic the complex TME as well as the restricted penetration of drugs and effector cells to the tumor center typically observed *in situ*, a Panc89 3D spheroid model was employed. In comparison to 2D monolayer culture of Panc89 L1+ cells, apoptosis induction by L1-bsAB was noticeably weaker in 3D spheroids of parental Panc89 population and higher overall concentrations had to be applied to achieve relevant tumor cell apoptosis. This finding reflects the higher heterogeneity of L1CAM expression of the Panc89 parental population modelling the tumor heterogeneity observed in PDAC tissues.^21^ Moreover, our findings are in line with previously published studies that demonstrate reduced efficacy of cytostatic and targeted anti-cancer drugs in 3D culture systems.^30,31^ Underlying mechanisms described in literature are decreased cell proliferation, altered cell metabolism, gradients in nutrient availability, impaired drug penetration and altered expression of receptor and drug transporter proteins.^30,31^ Furthermore, the 3D context differentially impacted efficacy of L1-bsAB1 and L1-bsAB2. When applied at a concentration of 1 nM, L1-bsAB1 induced significantly elevated levels of apoptosis and release of cytokines, whereas these effects were less pronounced after L1-bsAB2 treatment. Again, these differences may be explained by the distinct L1-bsAB design targeting different CD3 epitopes thereby resulting in altered effector cell activation kinetics. These observations highlight the added value of employing 3D models for preclinical bsAB evaluation in addition to 2D monolayer cultures to gain insights into dosage required for effective tumor cell killing in a more complex tumor context.

The TME dictates PDAC’s aggressive biology and its cellular components can attenuate cytostatic drug efficacy.^32–34^ To account for the impact of two key cell populations of the TME, we also employed Panc89 3D spheroids enriched with PMF or macrophages for evaluation of L1-bsAB efficacy. Interestingly, neither addition of PMF nor M1-like or M2- like macrophages relevantly impacted the effect of L1-bsAB1 or L1-bsAB2. It can be speculated that the limited duration of co-culture was too short to observe putative efficacy-limiting effects, e.g. mediated by direct cell-cell-interactions, paracrine activity of signaling molecules or ECM deposition, the latter forming a physical barrier for drug penetration and creating high intratumoral pressure. In contrast to cell viability, cytokine levels were differentially regulated by addition of PMF or macrophages to the 3D PDAC cell model. In the presence of PMF, IL-6 and IL-17A levels were significantly reduced after L1-bsAB1 treatment at 1 nM in comparison to respective mono-cultured 3D spheroids highlighting their immunomodulatory role in PDAC.^35,36^ In line with our findings, pancreatic cancer-associated fibroblasts have been shown to prevent T cell function by upregulation of immune checkpoints and secretion of cytokines inhibiting T cell function.^35,36^ To this end, Dominquez et al. and Krishnamurty et al. described a subtype of LRRC15+ cancer-associated fibroblasts (CAFs) induced by TGF-β signaling that promote expression of immune checkpoint proteins and suppress expression of effector cytokines in tumor-infiltrating T cells.^37,38^ Additionally, secretion of CXCL12 by CAFs expressing the fibroblast activation protein (FAP) was shown to promote T cell evasion from the tumor site in a PDAC mouse model.^23^ In our 3D spheroid model, the presence of a high proportion of either M1- or M2- macrophages (tumor cell: macrophages 1:2) reduced the release of T cell effector molecules IFN-γ, Granzyme B and IL-2 underscoring the previously described role of macrophages in impeding T cell effector function e.g. by expressing the immune checkpoint protein PD-L1.^39,40^

While (co-culture) spheroids model several aspects of tumor biology not represented by monolayer cultures, they lack the complexity of the TME of PDAC tissues. Moreover, the 2D and 3D tumor cell models described above and most experimental *in vitro* and *in vivo* data published to date on preclinical evaluation of bispecific TCE rely on the use of allogeneic effector cells. Therefore, unspecific allogeneic immune responses may influence results obtained on efficacy, T cell activation and cytokine release. Additionally, effector cells are often applied at high effector to target ratios in experimental settings unrepresentative of the heterogeneous infiltration of T cells into PDAC tissues resulting in defined areas with very limited T cell presence.^5^

To address these limitations, we evaluated L1-bsAB in OTSCs, a 3D tumor model derived from surgically resected PDAC tissue representing the preserved TME composition.^41^ Previous investigation confirmed that PDAC OTSCs mostly maintain cellular architecture and the expression of immunologic proteins for up to 92 h after initiation of culture.^42,43^ On a transcriptomic level, *ex vivo* cultivation of PDAC OTSCs resulted in very limited alteration of gene expression, supporting the suitability of OTSCs as a model to assess anti-tumor effects by (immune)therapeutics.^44,45^ Moreover, functional immunotherapy studies in human PDAC OTSCs demonstrated that resident intratumoral CD8-positive T cells can be reactivated *ex vivo* and mediate tumor-cell apoptosis, supporting the use of OTSCs to evaluate T-cell-dependent therapeutic strategies.^46^

Treatment with L1-bsAB2 resulted in induction of apoptosis in the OTSCs indicated by a significant increase in staining intensity for cleaved caspase 3 and a non-significant increase in ccK18 level in the culture supernatant even though overall L1CAM expression of tumor cells in the investigated OTSCs was rather low. L1-bsAB2-mediated apoptosis induction was accompanied by a trend towards reduced CK7 positive tumor cells and a reduced number of Ki67-positive proliferating tumor cells. Furthermore, increased release of INF-γ, Granzyme B and IL-2 into the supernatants was observed indicating T cell activation. Overall, this data indicates that L1-bsAB2 induces anti-tumor activity in OTSC representing a clinically-relevant experimental tumor model comprising an intact TME architecture and relying on autologous tumor infiltrating T cells as effector cells that had been present in the tumor stroma.

In conclusion, we designed two novel T cell-engaging L1-bsAB, both of which exert potent L1CAM-specific anti-tumor effects in different preclinical PDAC models considering the complex TME as well as the altered functional state of immune effector cells of PDAC patients. The findings further support the anti-tumor efficacy of L1-bsAB against PDAC cells under typical immunosuppressed TME conditions, as L1-bsAB2 was also effective in OTSCs, a model system representing the complete TME architecture present *in situ* and relying on autologous tumor infiltrating T cells as immune effector cells.

## Materials and Methods

### Cell lines and cell culture

Panc1 cells derived from a primary PDAC specimen were purchased from ATCC (Manassas, VA, USA). Panc1 L1+ and Panc1 L1- cell variants were generated by stable transfection with the pLVX vector containing sh-RNA targeting firefly luciferase (Panc1 L1+) or L1CAM (Panc1 L1-) as previously described.^12^ Panc89 cells derived from a PDAC lymph node metastasis were kindly provided by Prof. T. Okabe (University of Tokyo, Tokyo, Japan). Panc89 L1+ and L1- cells were obtained through single cell cloning. Both cell lines have been extensively characterized regarding L1CAM+ expression.^21,47^

All PDAC cell lines were cultured in Panc-medium (RPMI 1640 (Biochrom, Berlin, Germany) supplemented with 10% FCS, 1% L-glutamine and 1 % sodium pyruvate (PAN-Biotech, Aidenbach, Germany). Panc1 L+ and L1- cells were cultured in Panc-medium supplemented with 0.67 µg/ml puromycin (Invivo Gen, San Diego, CA, USA) for maintenance culture.

Absence of mycoplasma infection was regularly confirmed by testing for presence of mycoplasmatic enzymes (Lonza, Basel, Switzerland). Cell line authentification was performed by STR profiling.

### Pancreatic myofibroblasts

hPSC-eGFP cells, used as model for PMF, were isolated from a male chronic pancreatitis patient, immortalized using SV40 large T antigen and human telomerase (hTERT) and further transduced with an eGFP encoding plasmid (kindly provided by Matthias Löhr (Karolinska Institutet, Stockholm, Sweden).^48^ hPSC-eGFP cells were cultivated in hPSC-medium (DMEM high glucose (Biochrom, Berlin, Germany), 10% FCS (Pan-Biotech), 1 % L-glutamine (Biochrom), 1 µg/ml puromycin (Invivo Gen, San Diego, CA, USA)).

### Isolation of human PBMC from healthy donors and PDAC patients

Peripheral blood mononuclear cells (PBMC) of healthy donors were isolated from leucoreduction system chambers obtained during blood donation and provided by the Institute of Transfusion Medicine, UKSH Campus Kiel. PBMC from PDAC patients were isolated from EDTA-blood. Written informed consent was obtained from all healthy donors and patients. Research was approved by the ethics committee of the Medical Faculty of Kiel University and the University Hospital Schleswig-Holstein, Campus Kiel (Reference numbers: D601/25(A110/99), D490/17). PBMC were isolated using a Pancoll (PAN-Biotech) density gradient centrifugation protocol (800 xg, 25 min, room temperature (RT)). After centrifugation, the PBMC-layer was carefully collected and directly processed for experiments or cryostored in liquid nitrogen until thawed for experiments.

### Isolation and activation of human CD8+ T cells

First, to reduce monocyte populations in PBMC, 125×10^6^ PBMC were suspended in 10 ml RPMI 1640 supplemented with 1% FCS and incubated for 45 min in a 75 cm² flask. Then, the supernatant containing non-adherent cells (lymphocytes) was collected for further processing. The CD8+ cell population was purified via magnetic cell sorting (MACS) using a negative selection strategy (CD8+ T Cell Isolation Kit, human, Miltenyi Biotec, Bergisch Gladbach, Germany) to obtain untouched CD8+ T cells. The manufacturer’s instructions were modified using reduced antibody and bead concentrations (50%) and an extended incubation period (1.5 fold) as previously published.^49^ To activate CD8+ T cells, a 24-well plate was coated for 2 h at 37°C with 1.5 µg/ml anti-CD3 antibody (clone: OKT3, BioLegend, San Diego, USA). This was followed by two washing steps of the coated wells with PBS. Then 1.5×10^6^ CD8+ T cells were seeded per well in T cell medium (TCM) supplemented with 1.5 µg/ml anti-CD28 antibody (clone: CD28.2, BioLegend) and 60 ng/ml IL-2 (BioLegend). CD8+ T cells were cultured for 4 days prior to start of co-culture experiments.

### Generation of L1-bsAB

The DNA sequences of L1-bsAB1 and L1-bsAB2 were synthesized and cloned into a proprietary mammalian vector. Suspension adapted CHO-K1 (Evitria. Zürich-Fahrweid, Switzerland) cells were grown in chemically defined, animal component-free transfection medium and transfected with the expression vectors using a proprietary transfection reagent. Supernatant was harvested and filtered through a 0.2 μm filter, followed by purification of the bispecific antibodies by protein A affinity chromatography. L1-bsAB1 was purified using Mab Select SuRe (Cytiva, Freiburg, Germany). Analytical size exclusion chromatography (SEC) of purified L1-bsAB1 by MAbPac SEC (Thermo Fisher Scientific, Waltham, MA, USA) showed a monomer content of 100%. L1-bsAB2 was purified using Hi-Trap protein A HP (GE Healthcare, Chicago, IL, USA) followed by preparative size exclusion on a Superdex 200 Increase 10/300 G column (Cytiva). Monomericity of purified L1-bsAB2 was determined by size exclusion chromatography to be 100%. Both L1-bsAB1 and L1-bsAB2 were stored at 4°C in PBS pH 7.4 containing 100 mM arginine.

### Determination of binding affinity of L1-bsABs to L1CAM

Monovalent kinetic binding constants of L1-bsAB1 and L1-bsAB2 towards human L1CAM were determined by surface plasmon resonance using a Biacore T200 instrument (Cytiva). Bispecific antibodies were captured via their Fc regions on a C1 Chip (Cytiva) via immobilized Protein A, and soluble His-tagged human L1CAM was injected as analyte at a single concentration of 40 nM, using 10 mM 25 HEPES / 150 mM NaCl / 3 mM EDTA / 0.05 % Tween 20 as running buffer. Detected resonance units (sensorgrams) were fitted to a 1:1 Langmuir binding model and association (k_a_) and dissociation (k_d_) rate constants as well as the affinity constants (K_D_) were calculated.

### Fc receptor binding by L1-bsAB

Monovalent binding kinetics of L1-bsAB1 and L1-bsAB2 towards human neonatal Fc receptor (FcRn) and the Fcγ receptor IIIa isoforms F158 (FcγRIIIaF158) and V158 (FcγRIIIaV158) were captured by surface plasmon resonance. Details of the experimental approach are proprietary. Rituximab was used as a positive control. Binding affinity to FcRn was determined using a steady-state binding evaluation approach. The equilibrium resonance units (RU) measured at each analyte concentration were fitted using a four-parameter logistic (4PL) model to determine the apparent binding affinity. Absolute binding levels to FcγRIIIaF158 and FcγRIIIaV158 were expressed as resonance units, normalized to the FcγRIIIa capture level, and corrected for the mass of the analyte (L1-bsAB: 200 kDa, Rituximab (Roche, Basel, Switzerland): 145 kDa). Percent binding was calculated by dividing the absolute binding level observed for one-of the L1-bsAB by the absolute binding level observed for Rituximab, multiplied by 100.

### C1q binding by L1-bsAB

To determine C1q binding serial dilutions of L1-bsAB1, L1-bsAB2, and Rituximab (Roche), ranging from 0.02 to 20 µg/mL were coated on ELISA 96-well plates (Nunc Maxisorp, ThermoFisher Scientific, Roskilde, Denmark) in coating buffer (0.05 M sodium carbonate buffer, pH 9). The plates were blocked with 200 µl/well of ELISA diluent (0.1 M NaPO_4_ / 0.1 M NaCl / 0.1% gelatin / 0.05% Tween 20 / 0.05% ProClin300 (Sigma-Aldrich, Saint-Louis, Missouri, USA) for 1 h, and incubated for 2 h with 100 µl/well of 12 mg/ml human C1q (Quidel, San Diego, California, USA) in ELISA diluent. For detection of bound C1q, HRP-conjugated sheep anti-human C1q antibody (Biodesign) in ELISA diluent was added and incubated for 1 h. To control for coating efficiency of the different coated antibodies, HRP-conjugated goat anti-human Fc specific antibody (Jackson ImmunoResearch, West Grove, PA, USA) was added on a duplicate plate in ELISA diluent and incubated for 1h. Plates were washed after each incubation step with PBS/0.05% Tween 20, pH 7.4. The plates were finally developed with 100 µl of substrate buffer (PBS/0.012% H_2_O_2_) containing o-phenylenediamine dihydrochloride (Sigma Aldrich). The reaction was stopped by the addition of 100 ml of s4.5N H_2_SO_4_, and optical density (OD) was measured at 492 nm. To correct for background, the OD at 405 nm was subtracted from the OD at 492 nm. The corrected OD values were fitted using a four-parameter logistic (4PL) model. %binding was calculated by dividing the top plateau of the 4PL fit obtained with the bispecific antibodies by the top plateau obtained with Rituximab, multiplied by 100.

### T cell binding by L1-bsAB

The ability of the bispecific antibodies to engage CD3 on T cells was assessed by flow cytometry. Cryoconserved PBMC from one donor were thawed using CTL-anti-aggregate wash (C.T.L., Cleveland, Ohio, USA) and T cells were isolated using the EasySep™ Human T Cell Isolation Kit (STEMCELL Technologies, Vancouver, Canada) according to manufacturer’s instructions. T cells were counted and resuspended at 2×10^6^ cells/mL in cold FACS buffer (DPBS + 0.1%BSA) and 50 μL of this suspension were added to 96 well V bottom plates. Serial 5-fold dilutions of the bispecific antibodies ranging from 20 nM to 1.28 pM were prepared in cold FACS buffer and 50 µL of each dilution were added to the T cells. After an incubation of 1 h at 2-8°C, the cells were washed and resuspended in FACS buffer containing 1:100 diluted secondary anti-human IgG (R-Phycoerythrin-AffiniPure F(ab)_2_ Fragment Donkey Anti-human IgG (H+L)) (Jackson ImmunoResearch). Cells were incubated for another 30 min at 2-8°C, washed again, and resuspended in 25 µL cold FACS buffer. Seventyfive µL of fixation buffer (4% paraformaldehyde (PFA) in PBS) were then added to cells followed by an incubation at RT for 20 min. After an additional wash step cells were resuspended in 200 µL FACS buffer and acquired by an Acea Quanteon 4025 flow cytometer (Agilent, Santa Clara, CA, USA). A viability dye (Fixable viability dye eFluor 780, Thermo Fisher Scientific) was employed in parallel to exclude non-viable cells, and human IgG isotype control and untreated cells were employed to identify non-specific binding.

### Measurement of T cell mediated cytotoxicity

One hundred µL of a Panc-1 cell suspension in DMEM high glucose and L-glutamine medium (Scientific Laboratory Supplies, Nottingham, UK) were seeded at a density of 1×10^4^ cells/well into the wells of an xCELLigence E-plate 96 (Agilent). After in incubation for 30 min in the dark at room temperature to allow cells to settle to the bottom of the wells, the plate was transferred to the Real time cell analysis (RTCA) station (Agilent) inside a cell culture incubator, where incubation was continued for 24 h at 37°C/ 5% CO_2_ to allow cell attachment and proliferation. Impedance was recorded over 24 h as a measure of cell growth, using the xCELLigence RTCA system. Data acquisition was then paused, the plate was removed from the RTCA station, 50 µL of medium were removed, and 50 µL of a 4-fold concentrated dilution series of L1-bs-AB1 or L1-bsAB2 (10-fold dilutions, ranging from 10’000 pM to 1 pM) in X-VIVO 15 / 5%FBS (Lonza / Thermo Fisher) were added. Fifty µL of PBMC from a single donor (6 × 10^5^ cells / mL in X-VIVO 15 / 5% FBS), resulting in a final seeding density of 3×10^4^ cells and an effector: target ratio of 3:1. As controls, Panc-1 cells, PBMC, and Panc-1 cells with PBMC were incubated in the absence of bispecific antibodies. The plate was loaded back into the RTCA station and impedance measurement was continued for 72 h to monitor T cell mediated killing of Panc-1 cells. Raw data were expressed as Cell index and analyzed using the xCELLigence Immunotherapy Software and GraphPad Prism.

### Automated Fluorescence-microscopy assay for assessing L1-bsAB efficacy in 2D co-cultures of PDAC cell lines and PBMC

To identify PDAC cells in 2D co-cultures with effector cells, PDAC cells were first labelled with CellTracker green CMFDA (Thermo Scientific, Schwerte, Germany) following the manufacturer’s instruction and seeded at desired cell numbers in 200 µl Panc medium per well in transparent 96-well plates (Thermo Scientific). PDAC cells were allowed to adhere for 24 h prior to the start of co-culture and drug treatment. The next day, effector cells (PBMC or preactivated CD8+ T cells) were counted and added at the desired target: effector ratio (10:1) in 180 µl of TCM after removal of Panc medium from wells. For additional drug treatments, compounds were prediluted in a drug plate at 10x concentration. Then, 20 µl of phosphate buffered saline (PBS) (PAN-Biotech) as a negative control or antibodies (ctrl-bsAB, L1-bsAB1 or L1-bsAB2) were added to achieve the final drug concentration. After the treatment period of 8h for experiments with pre-activated CD8+ T cells and 48 h with PBMC, medium containing non-adherent cells was removed and centrifuged at 15000xg at 4°C. Supernatants were collected and stored at −80°C until further processing. Cell pellets were stained for flow cytometry analyses. Adherent cells were stained with Hoechst 33342 (1:5000 in PBS) (Sigma-Aldrich, München, Germany) for nuclei staining and Propidium Iodide (PI) (Sigma Aldrich, St. Louis, MO, USA) (1:50 in PBS) for detection of dead cells for 30 min. Then, cells were imaged with the NYONE® Scientific Imager (SYNENTEC GmbH, Elmshorn, Germany) recording bright field images and fluorescence signals. Data was processed with the YT-SOFTWARE® (SYNENTEC) using the virtual Cytoplasm (2F) operator to distinguish live tumor cells (Hoechst and Cell tracker green positive) from dead tumor cells (Hoechst, Cell tracker green and PI positive) and from remaining dead (Hoechst and PI positive) or live effector cells (Hoechst positive only). Viable tumor cells were automatically counted and counts were further processed by normalization to PBS control.

### Immunocytochemical staining of PDAC cell lines

Immunocytochemical L1CAM staining was performed to detect L1CAM protein levels in PDAC cell lines. First, 1 × 10^5^ cells per well of a 12-well flat bottom cell culture plate were seeded on glass coverslips in 2 ml of respective cell culture medium. After 24 h, coverslips with cells were washed for 5 min. All washing steps were performed with PBS (PAA). Cells were fixed for 15 min with 1 ml 4% (w/v) PFA (Thermo Fisher Scientific) followed by washing 3 x 5 min. Coverslips were incubated for 10 min in 0.3% hydrogen peroxide (H_2_O_2_) (Carl Roth, Karlsruhe, Germany) in ice-cold methanol (Th. Greyer, Renningen, Germany) to block endogenous peroxidase activity, followed by washing for 3 x 5 min. Incubation with 4% (w/v) bovine serum albumin (BSA) (Biomol, Hamburg, Germany) in PBS was performed for 20 min at RT to block unspecific binding. Then, coverslips were incubated for 45 min with the primary mouse IgG2a anti-human L1CAM antibody (clone 9.3) or its respective isotype-matched control antibody (both kindly provided by Prof. Dr. Peter Altevogt, DKFZ Heidelberg, Germany) at a final concentration of 10 µg/ml in a humidified chamber at RT. After washing, coverslips were incubated with EnVision+ System-HRP Labelled Polymer Anti-mouse (Agilent) for 30 min followed by repeated washing steps. Aminoethyl carbazole (AEC) (Agilent) was used for the chromogenic reaction. Nuclei were stained for 2 −5 min with Mayer’s Haemalaun (Sigma-Aldrich, München, Germany) and subsequently rinsed with water. Coverslips were conserved with aqueous-based glycerol gelatine (Merck, Darmstadt, Germany) on microscope glass slides. Images were acquired with the Lionheart FX Automated Microscope (Biotek, Bad Friedrichshall, Germany).

### Multiplex Cytokine Assay

Levels of cytokines and T cell effector molecules were analyzed in supernatants from 2D and 3D co-cultures as well as from OTSC using the LEGENDplex^TM^ Human CD8/NK V2 Panel omitting analyses of IL-4, TNF-alpha, sFAS and sFASL for evaluation of 2D experiments. Measurements were performed on a BD FACSymphony^TM^ A1 flow cytometer (BD Bioscience, Franklin Lakes, NJ, USA) and analyzed with the LEGENDplex^TM^ data analysis software (BioLegend).

### ccK18 Enzyme-Linked Immunosorbent Assay (ELISA)

Levels of the soluble M30 neoepitope released upon caspase-cleavage of cytokeratin-18 (ccK18) were determined by M30 CytoDeath**^TM^** ELISA (PEVIVA, Diapharma, West Chester, PA, USA) in culture supernatants allowing specific detection of epithelial/PDAC cell apoptosis. The assay was performed according to manufacturer instruction. The readout was performed on a TECAN Infinite® 200 PRO microplate reader (TECAN, Männedorf, Switzerland).

### Flow Cytometry

Immunofluorescence staining and flow cytometry were performed to characterize T cell and PBMC populations of healthy donors and PDAC patients. First, cells were washed with PBS and incubated for 10 min in 5% (v/v) human Fc receptor blocking reagent (Miltenyi Biotec GmbH, Bergisch Gladbach, Germany) in FACS buffer (0.5% (w/v) BSA, 2 mM EDTA in PBS) and subsequently washed. Immunofluorescence staining was performed with fluorophore-labeled specific primary antibodies (AB) or the respective isotype control in FACS buffer for 30 min in the dark at 4°C. The following specific primary mouse anti-human AB were used: anti-CD4-APC (clone RPA-T4, 2.5 µg/ml), anti-CD8a-FITC (clone HIT8a, 10 µg/ml), anti-PD-1-PE (clone A17188B, 1.25 µg/ml), anti-PD-L1-PECy7 (clone MIH3 10 µg/ml), anti-CD69-PECy7 (clone FN50, 5 µg/ml), anti-CD25-APC (clone BC96, 5 µg/ml). Isotype control antibodies IgG1-FITC, -PECy7 or -APC (all clone MOPC-21) and IgG2b-PE (clone MPC-11) were used at the same concentration as the specific primary AB (all Biolegend). Next, samples were washed twice and fixated with 1% (v/v) PFA in FACS buffer. All washing steps were performed in a total volume of 200 µl MACS buffer in a 96-well V-bottom plate with subsequent centrifugation for 8 min at 300· g and 4°C. Flow cytometric measurements were performed on a MacsQuant X (Miltenyi Biotec GmbH, Bergisch Gladbach, Germany). For each measurement, signals from at least 2×10^4^ lymphocytes were recorded. FlowJo v10.8.1 (BD Bioscience, Franklin Lakes, New Jersey, USA) was used for data analysis.

To quantify the relative levels of surface molecules, the intensity of specific immunofluorescence staining was normalized to the staining intensity of the respective isotype control antibody calculating the ratio of median fluorescence intensities (MFI ratio = MFI (specific staining)/ MFI (respective isotype control staining)).

### Monocyte isolation and generation of macrophages

CD14+ monocytes were isolated by negative magnetic cell separation (BioLegend, Fell, Germany) from PBMC obtained via density gradient centrifugation from leukoreduction system chambers of healthy blood as described above. A modified manufacturer’s protocol was applied for magnetic cell separation using reduced antibody and bead concentrations (25% of original concentration) and increased incubation times (15 min per incubation step). After isolation, CD14+ monocytes were differentiated into M1- and M2- macrophages. To this end, isolated monocytes were counted, centrifuged and resuspended in either M1-medium (RPMI-1640, 5% FCS, 1% penicillin/streptomycin (P/S) and 1% L-glutamine) or M2- medium (RPMI-1640, 1% FCS and 1% P/S). For differentiation, 10 monocytes were seeded into petri dishes and stimulated with either 2,4 ng/ml GM-CSF (BioLegend) for M1-macrophages or 50 ng/ml M-CSF (BioLegend) for M2- macrophages for 5-7 days.

### 3D PDAC cell spheroids in the absence or presence of different stromal cells

To achieve spheroid formation, cells were seeded in 96-well ultra-low attachment (ULA) plates (faCellitate, Mannheim, Germany) in 200 µl medium for 24 h. Spheroids were either generated from PDAC cells only (mono), from co-cultures of PDAC cells with macrophages differentiated into either M1- or M2- phenotype at two different PDAC: macrophage ratios (1:1 and 1:2) or with PMF at two ratios (1:1 and 1:2). For co-cultures at 1:1 ratio, 1×10 cells of each type were seeded, while for the 1:2 ratio, 0,667×10 PDAC cells and 1,333×10 macrophages or PMF were seeded. The number of PDAC cells seeded for formation of mono-cultured PDAC cell spheroids was adjusted to the respective number of PDAC cells seeded for either co-culture. Successful spheroid formation was regularly confirmed by light microscopy with the NYONE® Scientific Imager (SYNENTEC GmbH). After spheroid formation, 180 µl of medium were replaced by 130 µl TCM containing 2×10 PBMC (from same donor previously used for monocyte isolation) as effector cells to achieve an effector: target ratio of 10:1. Next, spheroids were treated with ctrl-bsAB or either L1-bsAB at final concentrations of 100 pM and 1 nM for 48 h. To account for potential unspecific tumor cell killing by PMBC, an untreated control condition was included that underwent medium exchange only. After treatment, the supernatant of three matching wells (technical replicates) was pooled, centrifuged at 300xg at RT for 5 min and supernatants were cryopreserved at −80 °C for further analyses.

### Organotypic slice cultures (OTSCs) from resected PDAC specimen

OTSC from four treatment-naïve patients who underwent PDAC resection were established as previously published.^50^ Research was approved by the ethics committee of the University of Lübeck and the University Hospital Schleswig-Holstein, Campus Lübeck (Reference number: #16-281) Clinical characteristics of patients are described in **Table 1**. In brief, resected tissue was pathologically evaluated and adjusted to approximately 0.5×0.5×0.5 cm using a sterile scalpel. After adjustment the tissue was stabilized in 8 % low-melt agarose (Roth Chemie, Karlsruhe, Germany) and further processed by a vibratome Leica VT1200 S (Leica Biosystems, Nussloch, Germany). OTSC were transferred onto semiporous hydrophilic polytetrafluoroethylene-ethylene (PTFE) tissue culture inserts (Milipore, Darmstadt, Germany) (pore size 0.4 µm). These PTFE inserts were placed in a 24-well plate containing 400 µl of advanced DMEM/F12 medium supplemented with 10 % FBS, 1 % penicillin/streptomycin (all Thermo Fisher Scientific) and incubated at 37 °C and 5 % CO_2_ in a humified atmosphere for 72 h. Cultivation medium was changed every 24 h. Treatment with respective drugs was initiated after 24 h of equilibration. Drug treatment was performed for 48 h.

### Immunohistochemical staining of OTSC

After treatment, OTSC were fixed in 4.5 % formalin and embedded in paraffin followed by hematoxylin-eosin (H&E) staining and IHC staining. For this purpose, OTSC were sectioned horizontally (4 µm) using a microtome (Rotary Microtome HM355S, Thermo Fisher Scientific). Sections were deparaffinized in xylene and rehydrated in graded ethanol baths followed by antigen retrieval with antigen retrieval pH 9.0 at sub-boiling temperature for 20 min. Followed by three washing steps with PBS each for 5 min at RT. To reduce non-specific protein interactions, blocking was performed using 4 % BSA/PBS for 20 min followed by primary antibody (Mouse anti-human L1CAM (R&D Systems #MAM7771-100, R&D Systems, Minneapolis, USA) over night at 4 °C. After three washing steps with PBS, sections were incubated with Envision anti-mouse HRP (Agilent) for 30 min at RT in a wet chamber, followed by three washing steps with PBS. AEC substrate (Agilent) solution was added for 20 min followed by another washing step (3x PBS). Finally, sections were stained with Mayer’s hemalum (Sigma-Aldrich, München, Germany) for 30 sec. Excess coloring solution was rinsed off under running water and sections were mounted with Kaiser’s Glycerin gelatin (Merck Millipore, Billerica, USA). Serial OTSC sections were additionally stained for CK7, Ki67 and CC3 to assess tumor cell content, proliferation and apoptosis, respectively. CK7 was detected using mouse anti-human CK7 clone OV-TL 12/30 at 1:200 (M7018; Dako, Düsseldorf, Germany), Ki67 using mouse anti-human Ki67 clone MIB-1 at 1:200 (M7240; Dako, Düsseldorf, Germany), and CC3 using rabbit anti-human cleaved caspase 3 clone 5A1E at 1:400 (9664; Cell Signaling Technology, Danvers, MA, USA). Detection was performed using biotinylated secondary antibodies, the VECTASTAIN ABC system (Newark, CA, USA) and DAB substrate, followed by hematoxylin counterstaining and Aquatex mounting (Merck, Darmstadt, Germany).

### PDAC patient characteristics

Blood samples from treatment-naïve patients (n=13) with PDAC were provided by the Translational Interdisciplinary Biobank Kiel (TRIBanK, Kiel, Germany). Written informed consent was obtained prior to blood sampling. Research was approved by the ethics committee of the Medical Faculty of Kiel University and the University Hospital Schleswig-Holstein, Campus Kiel (Reference numbers: D601/25(A110/99)). PBMC were isolated from blood samples obtained immediately before pancreatectomy or before start of chemotherapy in chemotherapy-naïve PDAC patients. Clinical and pathological data of PDAC patients are listed in **Table 1**. The TNM stage was determined pathologically for patients after tumor resection and a clinical stage is available for the patients with unresectable metastatic PDAC.

### Statistics

Statistical analysis was performed using GraphPad Prism Version 10.6.1 (GraphPad Software Inc., La Jolla, USA). Normality was tested by using the Shapiro-Wilk test. If the samples passed the normality test, Welch’s t-test was applied to compare two groups. Depending on the experimental design, either an ordinary one-way ANOVA or two-way ANOVA followed by Tukey’s multiple comparisons test was performed two compare more than two groups. Not-normally distributed data was analyzed by the Kruskal-Wallis test followed by Dunn’s multiple comparisons test. Normally distributed data is presented by column bar graph with mean and standard deviation, not normally distributed data is depicted by bar graphs with median and interquartile range in both directions. Results were considered as statistically significant for p-values < 0.05. Significance levels are indicated by asterisks: * = *p* < 0.05, **= *p* < 0.01, *** = *p* < 0.001, ****= *p* < 0.0001.

## Data Availability Statement

The data that support the findings of this study are available from the corresponding author upon reasonable request.

## Acknowledgments

We thank Sandra Schöne for excellent technical assistance. We would like to acknowledge the support of the CYTO KIEL | Cytometry Facility of the UKSH and we thank Anna Willms and Reinhild Geisen from SYNENTEC for excellent support regarding the use of their imagers. We thank the Translational Interdisciplinary Biobank Kiel (TRIBanK) for providing blood samples and clinical data for this study. The TRIBanK is a member of the PopGen 2.0 Biobanking Network (P2N) Kiel. We thank the Interdisziplinäres Centrum für Biobanking-Lübeck (ICB-L) for providing tissue samples and clinical data for this study. The authors AMW and RB gratefully acknowledge support from the Detlef Zillikens Clinician Scientist Academy of Precision Health in Schleswig-Holstein (PHSH) and the Clinician Scientist Academy Kiel. Figures depicting workflows were created using BioRender.com.

## Author contributions

**Anna Maxi Wandmacher:** Conzeptualization, Formal analysis, Funding acquisition, Investigation, Methodology, Visualization, Writing-original draft, Writing-review and editing. **Annika Brauer:** Formal analysis, Investigation, Methodology, Visualization, Writing-original draft, Writing-review and editing. **Charlotte Kayser:** Formal analysis, Investigation, Visualization, Writing-review and editing. **Caj Stach:** Investigation, Writing-review and editing. **Jakob Werner:** Investigation, Writing-review and editing. **Silje Beckinger:** Investigation, Methodology, Writing-review and editing. **Tina Daunke:** Investigation, Methodology, Writing-review and editing. **Benjamin Heckelmann:** Investigation, Methodology, Writing-review and editing. **Ashinikumar Hidam**: Investigation, Methodology, Writing-review and editing. **Olha Lapshyna:** Investigation, Methodology, Writing-review and editing. **Daniela Wesch:** Resources, Writing-review and editing**. Anne-Sophie Mehdorn**: Resources, Writing-review and editing**. Christoph Röcken:** Resources, Methodology, Writing-review and editing**. Rüdiger Braun:** Resources, Methodology, Writing-review and editing**. Flavio Mehli:** Investigation, Methodology, Conzeptualization, Writing-original draft, Writing-review and editing. **Anne Schmidt:** Methodology, Conzeptualization, Writing-review and editing. **Gunther Spohn**: Methodology, Conzeptualization, Writing-original draft, Writing-review and editing. **Susanne Sebens**: Conzeptualization, Funding acquisition, Investigation, Supervision, Project administration, Writing-original draft, Writing-review and editing.

