## Supplementary data for "L1CAMxCD3 bispecific antibodies exert potent anti-tumor effects in preclinical pancreatic cancer models considering the complex tumor stroma"

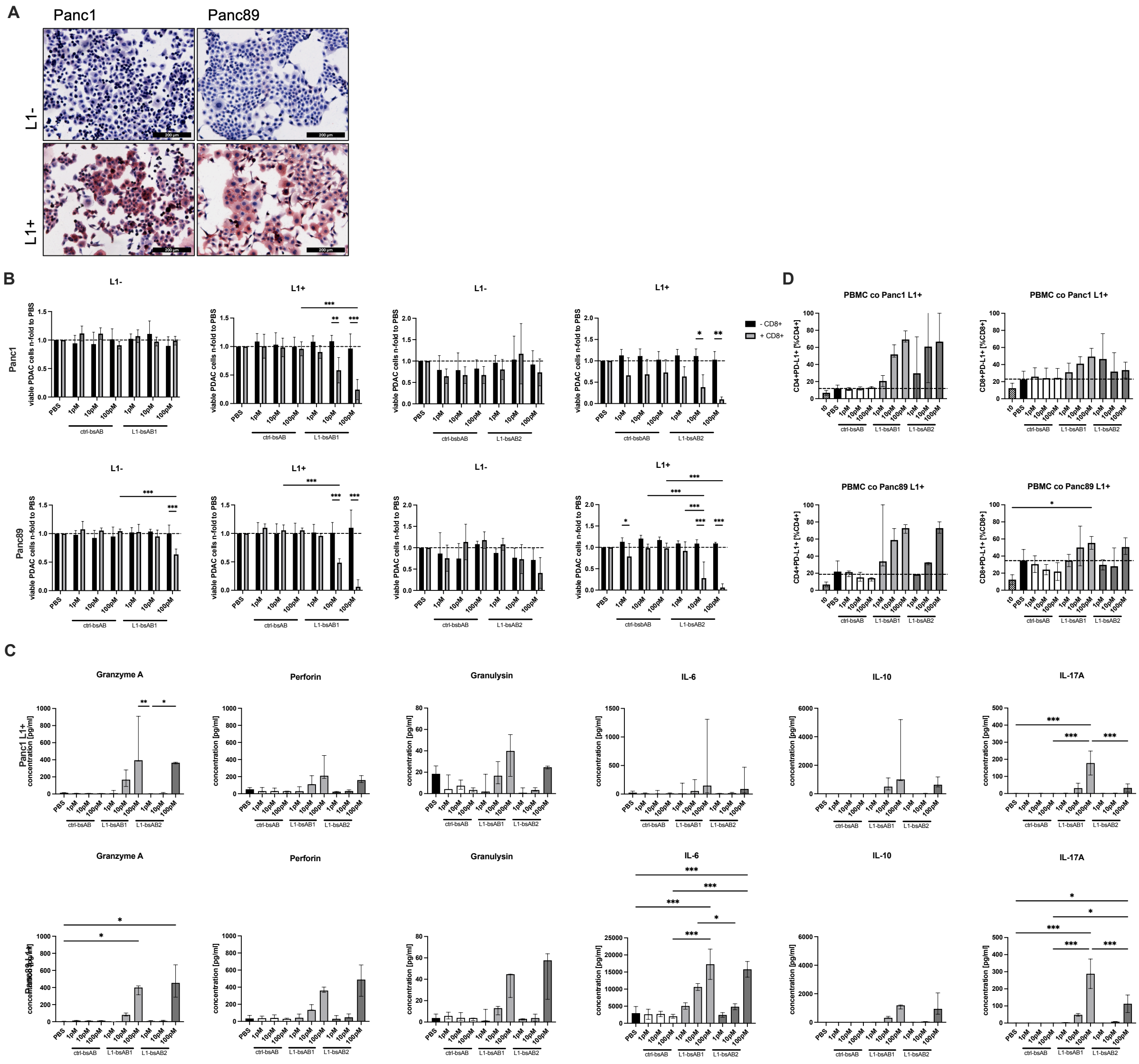


**Supplemental Figure 1: Bs-AB antibody efficacy in co-culture with pre-activated CD8+ T cells, release of cytokines from PBMC and expression of immunoregulatory receptor PD-L1 in T cells after co-culture with PDAC cell lines and treatment with L1-bsAB**

(**A**) Representative images of immunocytochemical staining of L1CAM in PDAC cell lines. Panc1 L1-, Panc1 L1+, Panc89 L1- and Panc89 L1+ were seeded on coverslips and stained for L1CAM expression (red) 24h after seeding. Nuclei were stained with Mayer's Hemalum (blue). Scale bar: 200 µm. (**B**) Anti-tumor efficacy of L1-bsAB was determined by automated cell imaging. PDAC cell lines with differential L1CAM expression were co-cultured with pre-activated CD8+ T cells from healthy donors (effector: target ratio 10:1) for indicated conditions and treated with PBS, ctrl-bsAB, L1-bsAB1 or L1-bsAB2 for 8 h. Live PDAC cells (Cell tracker green and Hoechst positive, PI negative) were counted with an automated cell imager after treatment. Data is presented as n-fold of viable cells treated with PBS. (**C**) Release of Granzyme A, Perforin, Granulysin, IL-6, IL-10 and IL-17A from co-cultured PBMC with Panc1 L1+ (upper panel) and Panc89 L1+ (lower panel) into the culture supernatant measured by multiplex assay. Supernatant was collected 48 h after treatment and initiation of co-culture. Data is presented as concentration [pg/ml]. (**D**) Flow cytometry analysis of PBMC 48h after initiation of drug treatment and co-culture with indicated PDAC cells. Staining was performed for immunoregulatory ligand PD-L1. Data is shown as frequency of PD-L1 positive cells (%) in the respective T cell population (CD4+ or CD8+). (**B-D**) Data is presented as mean (SD) for normally distributed data and median with interquartile range for not normally distributed data. Every analysis was performed with n=3 independent experiments using PBMC from 3 different donors and significances are indicated by asterisks: * = $p<0.05$; ** = $p<0.01$; *** = $p<0.001$.

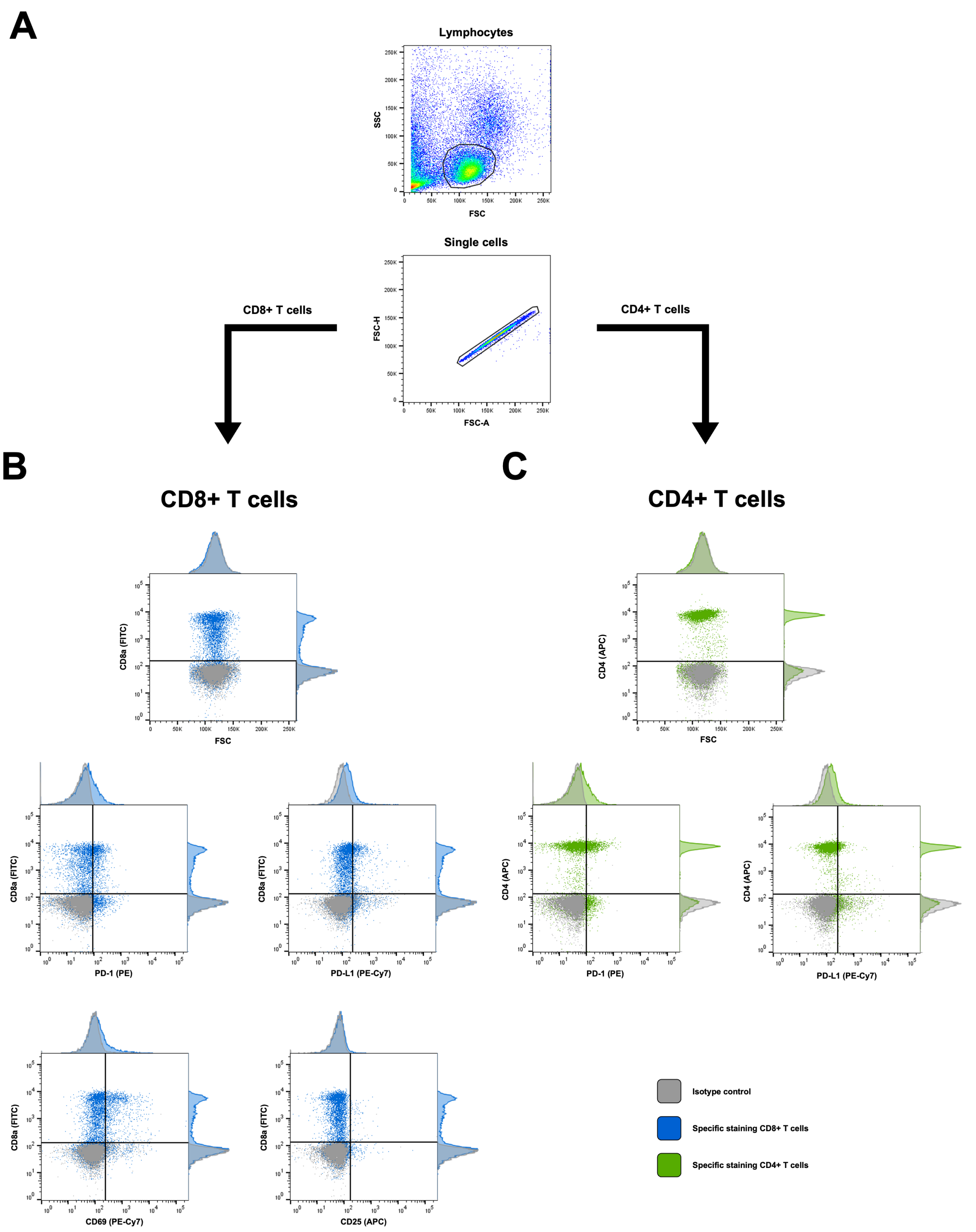


**Supplemental Figure 2: Gating Strategy for characterization of PBMC**

(**A**) Lymphocytes were identified as a distinct cluster based on SSC and FSC, next single cells were selected based on FSC-A and FSC-H. (**B**) Gates for CD8+, CD8+PD-1+, CD8+PD-L1+, CD8+CD69 and CD8+CD25+ (**C**) and for CD4+, CD4+PD-1+, CD4+PD-L1+ were defined based on negative isotype controls.

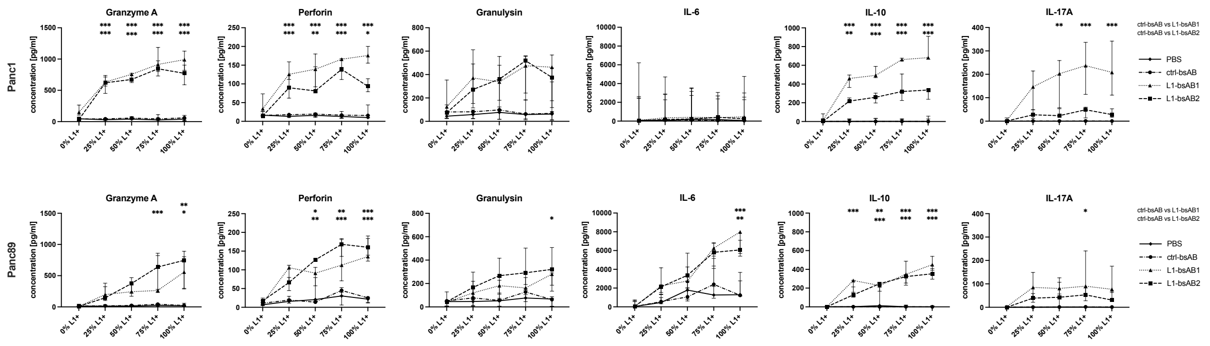


**Supplemental Figure 3: Release of cytokines from PBMC of healthy donors after co-culture with PDAC cell lines seeded at defined ratios of L1- and L1+ cells and treatment with L1-bsAB**

Release of Granzyme A, Perforin, Granulysin, IL-6, IL-10 and IL-17A from co-cultured PBMC (Effector: target ratio 10:1) with Panc1 (upper panel) and Panc89 (lower panel) into the culture supernatant measured by multiplex assay. Supernatant was collected 48 h after treatment (concentration of all indicated antibodies 100 pM) and initiation of co-culture. Data is presented as concentration [pg/ml]. Data from n=3 independent experiments using PBMC from 3 different donors is shown. Data is presented as mean (SD) for normally distributed data and median with interquartile range for not normally distributed data. Significances are indicated by asterisks: * = p<0.05; ** = p<0.01; *** = p<0.001.

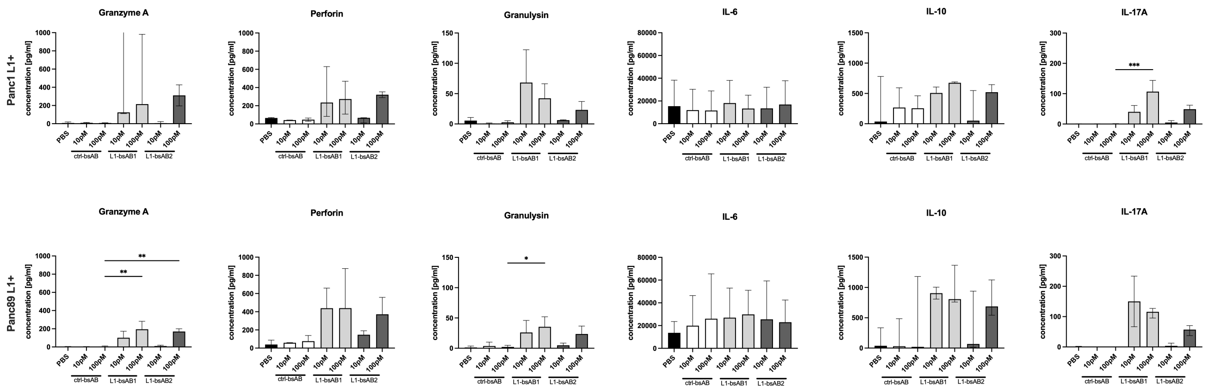


**Supplemental Figure 4: Release of cytokines from PBMC of PDAC patients after treatment with L1-bsAB and co-culture with PDAC cell lines**

Release of Granzyme A, Perforin, Granulysin, IL-6, IL-10 and IL-17A from PBMC from PDAC patients (Effector: target ratio 10:1) co-cultured with Panc1 L1+ (upper panel) and Panc89 L1+ (lower panel) into the culture supernatant measured by multiplex assay. Supernatant was collected 48 h after treatment and initiation of co-culture. Data is presented as concentration [pg/ml]. Data from n=3 independent experiments using PBMC from 3 different donors is shown. Data is presented as mean (SD) for normally distributed data and median with interquartile range for not normally distributed data. Significances are indicated by asterisks: * = p<0.05; ** = p<0.01; *** = p<0.001.

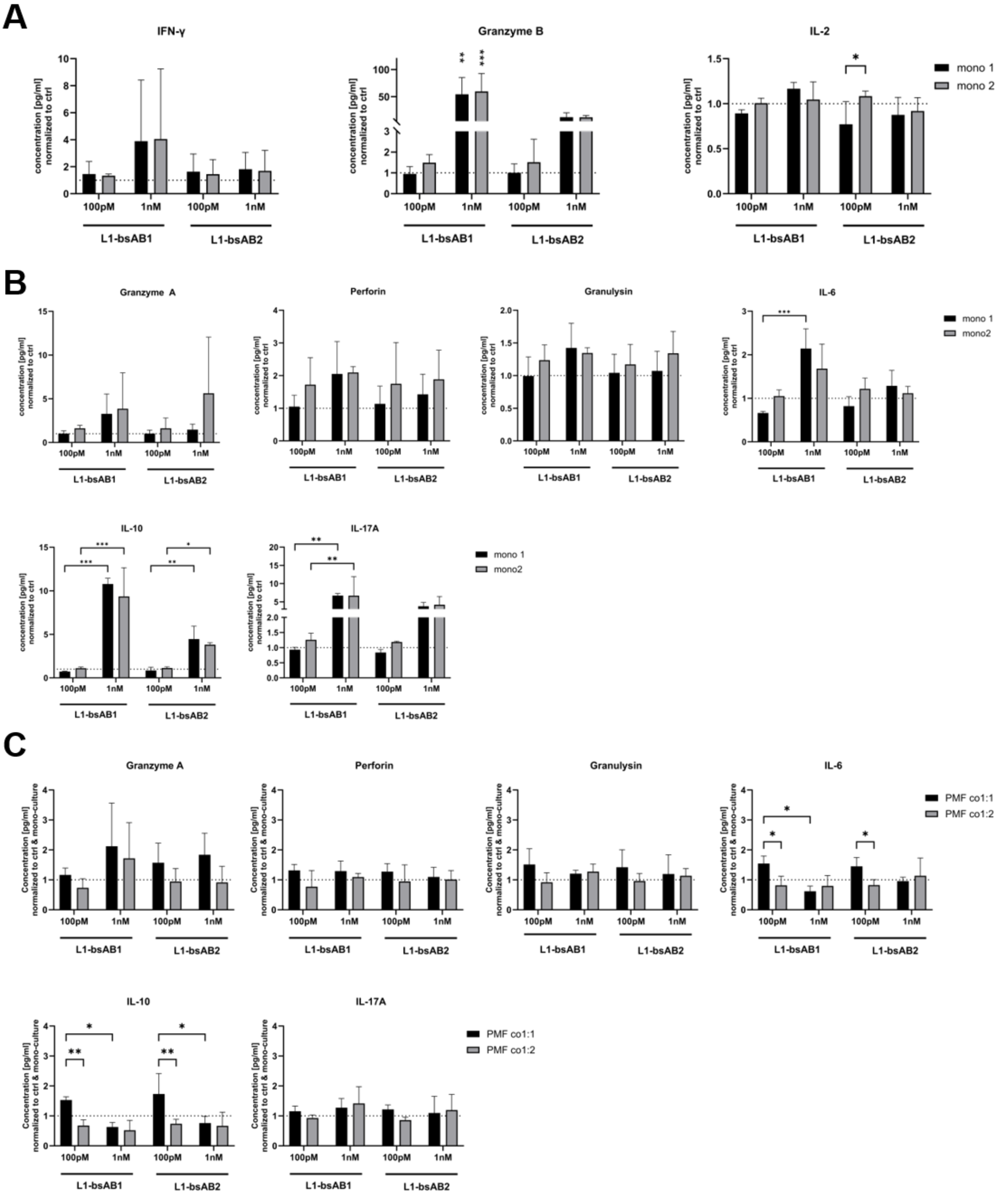


**Supplemental Figure 5**: **Release of cytokines from PBMC after co-culture with PDAC mono and PMF co-cultured spheroids and treatment with L1-bsAB**

Release of IFN-γ, Granzyme B, IL-2 **(A)**, Granzyme A, Perforin, Granulysin, IL-6, IL-10 and IL-17A from PBMC co-cultured with Panc89 parental **(B)** mono-culture spheroids and **(C)** co-culture spheroids with PMF after L1-bsAB treatment (Effector: target ratio 10:1), into the culture supernatant measured by multiplex assay. Supernatant was collected 48 h after treatment and initiation of PBMC co culture. Data from n=3 independent experiments using PBMC from 3 different donors is shown. Data is presented as mean (SD) for normally distributed data and median with interquartile range for not normally distributed data. Significances are indicated by asterisks: * = $p<0.05$; ** = $p<0.01$.

**A**

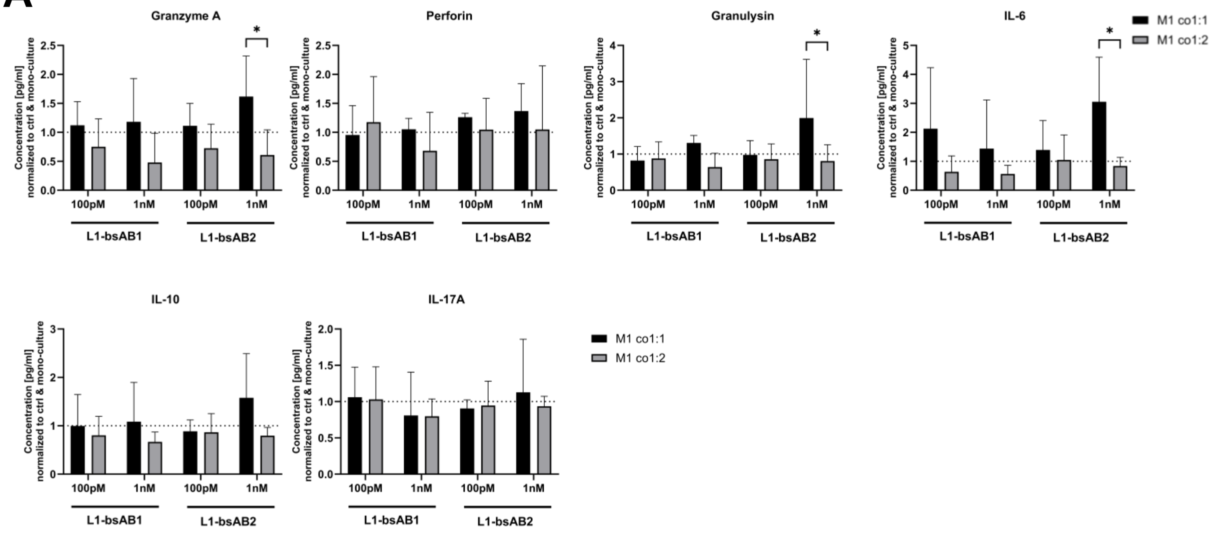


**B**

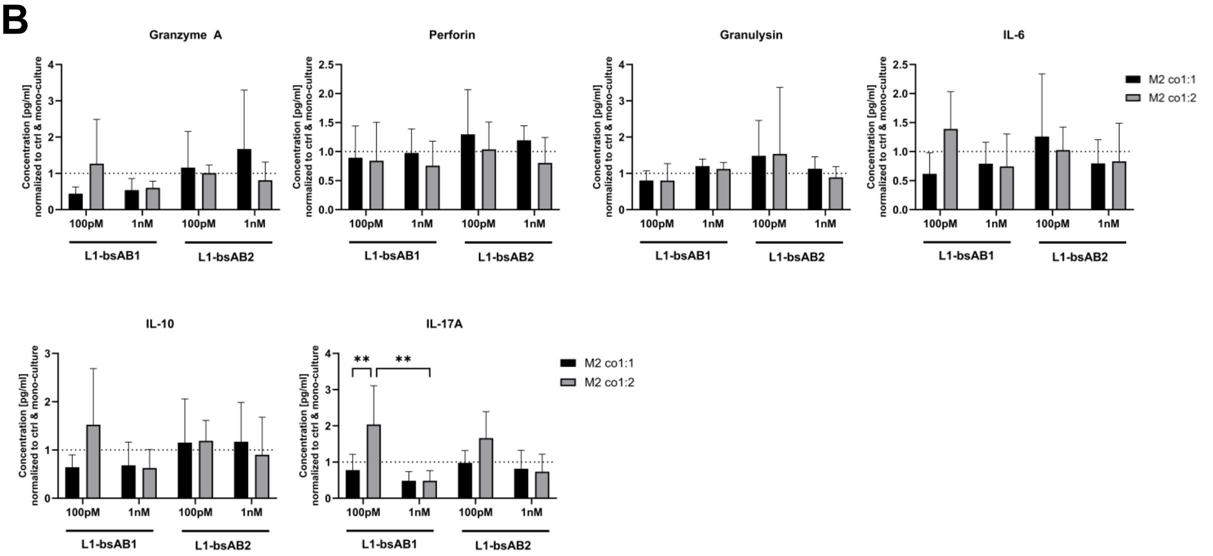


**Supplemental Figure 6**: **Release of cytokines from PBMC after co-culture with PDAC mono and PMF co-cultured spheroids and treatment with L1-bsAB**

Release of Granzyme A, Perforin, Granulysin, IL-6, IL-10 and IL-17A from PBMC, co-cultured with Panc89 parental co-culture spheroids with **(A)** M1-like macrophages or **(B)** M2-like macrophages after L1-bsAB treatment (Effector: target ratio 10:1), into the culture supernatant measured by multiplex assay. Supernatant was collected 48 h after treatment and initiation of PBMC co-culture. Data from n=4 independent experiments using PBMC from 4 different donors is shown. Data is presented as mean (SD) for normally distributed data and median with interquartile range for not normally distributed data. Significances are indicated by asterisks: * = p<0.05; ** = p<0.01.

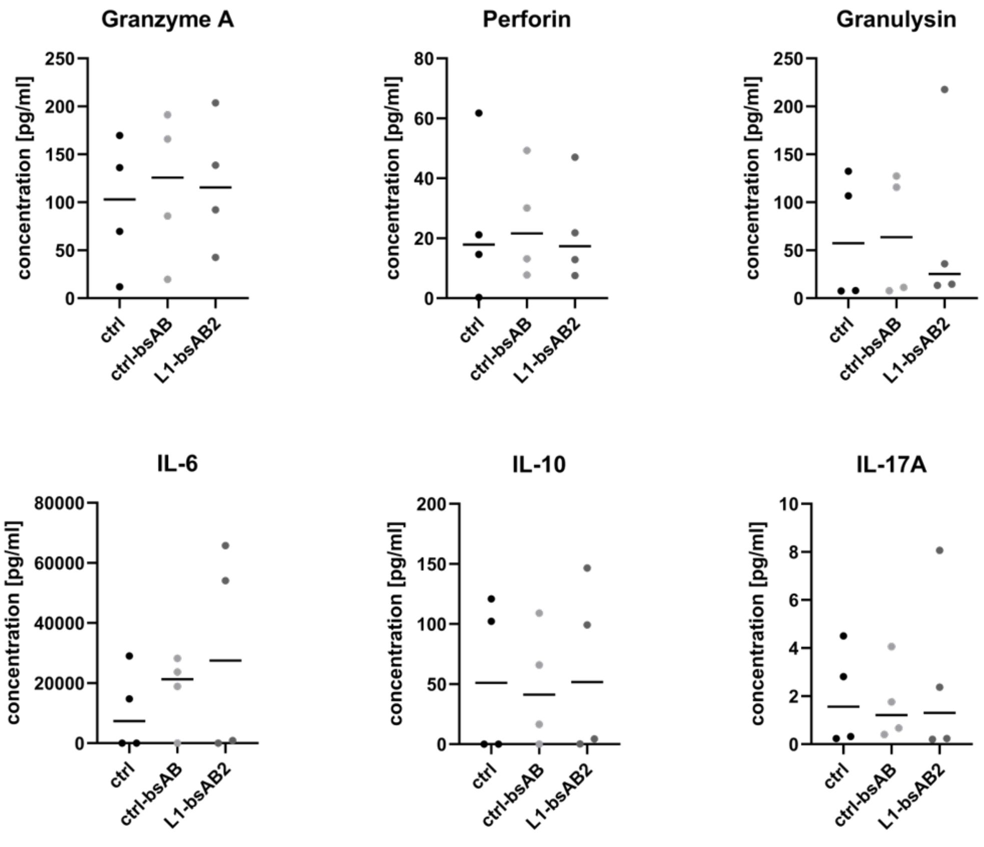


**Supplemental Figure 7**: **Release of cytokines after L1-bsAB treatment of OTSC**

Release of Granzyme A, Perforin, Granulysin, IL-6, IL-10 and IL-17A in the supernatants of OTSCs after L1-bsAB2 treatment for 48 h measured by multiplex assay. Data from n=4 independent experiments using OTSCs from 4 different PDAC patients is shown. Data is presented as mean for normally distributed data and median for not normally distributed data with data points representing individual OTSCs.
